# Epidermal PAR-6 and PKC-3 are essential for postembryonic development of *Caenorhabditis elegans* and control non-centrosomal microtubule organization

**DOI:** 10.1101/2020.07.23.217679

**Authors:** Victoria G. Castiglioni, Helena R. Pires, Rodrigo Rosas Bertolini, Amalia Riga, Jana Kerver, Mike Boxem

## Abstract

The cortical polarity regulators PAR-6, PKC-3 and PAR-3 are essential for the polarization of a broad variety of cell types in multicellular animals, from the first asymmetric division of the *C. elegans* zygote to apical–basal polarization of epithelial cells. In *C. elegans*, the roles of the PAR proteins in embryonic development have been extensively studied, yet little is known about their functions during larval development. Using auxin-inducible protein depletion, we here show that PAR-6 and PKC-3, but not PAR-3, are essential for postembryonic development. We also demonstrate that PAR-6 and PKC-3 are required in the epidermal epithelium to support animal growth and molting, and the proper timing and pattern of seam cell divisions. Finally, we uncovered a novel role for PAR-6 in controlling the organization of non-centrosomal microtubule arrays in the epidermis. PAR-6 was required for the localization of the microtubule organizer NOCA-1/Ninein, and microtubule defects in a *noca-1* mutant are highly similar to those caused by epidermal PAR-6 depletion. As NOCA-1 physically interacts with PAR-6, we propose that PAR-6 promotes non-centrosomal microtubule organization through localization of NOCA-1/Ninein.

**Summary:** Using inducible protein degradation, we show that PAR-6 and PKC-3/aPKC are essential for postembryonic development of *C. elegans* and control the organization of non-centrosomal microtubule bundles in the epidermis, likely through recruitment of NOCA-1/Ninein.

## Introduction

Polarity is a near universal property of cells that is essential for establishing proper cellular architecture and function. Epithelial cells – one of the major polarized animal cell types – polarize along an apical–basal axis and establish molecularly and functionally distinct apical, basal, and lateral membrane domains. The boundary between apical and lateral domains is marked by the presence of cell–cell junctions that provide adhesion between cells and prevent unwanted paracellular passage of molecules. The polarization of epithelial cells is orchestrated by conserved cortical polarity regulators that establish opposing membrane domains through mutually antagonistic interactions. In metazoans, the partitioning defective (PAR) proteins Par3, Par6, and atypical protein kinase C (aPKC) play a central role in the establishment of epithelial cell polarity. These highly conserved polarity regulators are essential determinants of apical domain identity, and are required for the positioning, maturation, and maintenance of apical cell junctions (Achilleos et al., 2010; Franz and Riechmann, 2010; Georgiou et al., 2008; Harris and Peifer, 2004; Harris and Peifer, 2005; Harris and Tepass, 2008; Hutterer et al., 2004; Izumi et al., 1998; Joberty et al., 2000; Leibfried et al., 2008; Lin et al., 2000; Totong et al., 2007; Wodarz et al., 2000; Yamanaka et al., 2001).

Par6 and Par3 are both PDZ domain containing scaffold proteins that can interact with each other, with aPKC, and with numerous other proteins. Par6 and aPKC form a stable subcomplex by interacting through their PB1 domains (Hirano et al., 2005; Wilson et al., 2003). The association of Par6–aPKC with Par3 is dynamic. In *C. elegans* zygotes, PAR-6/PKC-3 shuttle between a kinase inactive complex with PAR-3 that promotes anterior segregation, and an active complex with the small GTPase CDC-42 (Aceto et al., 2006; Beers and Kemphues, 2006; Rodriguez et al., 2017; Wang et al., 2017). In epithelia, Par3 can promote the apical recruitment of Par6–aPKC (Franz and Riechmann, 2010; Harris and Peifer, 2005; Hutterer et al., 2004; Joberty et al., 2000; Lin et al., 2000; Wodarz et al., 2000). In mature epithelia, however, the bulk of Par3 segregates to the apical/lateral border, where it plays an essential role in the positioning and assembly of apical junctions (Achilleos et al., 2010; Georgiou et al., 2008; Harris and Peifer, 2004; Harris and Tepass, 2008; Izumi et al., 1998; Leibfried et al., 2008; Totong et al., 2007; Yamanaka et al., 2001). The release of Par3 in epithelia depends on phosphorylation of Par3 by aPKC, and involves handoff of Par6–aPKC to Cdc42 and the epithelial specific Crumbs polarity complex (Bilder et al., 2003; Harris and Peifer, 2005; Hong et al., 2003; Krahn et al., 2010; Morais-de-Sá et al., 2010; Nagai-Tamai et al., 2002; Nunes de Almeida et al., 2019; Walther and Pichaud, 2010).

In addition to interactions that mediate the subcellular localization of Par6–aPKC or Par3, both Par6 and Par3 can interact with effector proteins to connect cortical polarity with downstream pathways (McCaffrey and Macara, 2009). For example, Par3 modulates phospholipid levels by recruiting the lipid phosphatase PTEN to cell junctions (Feng et al., 2008; Martin-Belmonte et al., 2007; Pinal et al., 2006; von Stein et al., 2005), inhibits Rac activity by binding to and inactivating the RacGEF Tiam1 (Chen and Macara, 2005; Mertens et al., 2005), and mediates spindle positioning in Drosophila neuroblasts through recruitment of Inscuteable (Schober et al., 1999; Wodarz et al., 1999). For Par6, fewer downstream targets have been described. In mammals, Par6 can recruit the E3 ubiquitin ligase Smurf1 to promote degradation of the small GTPase RhoA, causing dissolution of tight junctions (Ozdamar et al., 2005; Sánchez and Barnett, 2012; Wang et al., 2003). Par6 can also bind to the nucleotide exchange factor ECT2 to regulate epithelial polarization and control actin assembly at metaphase in dividing epithelial cells (Liu et al., 2004; Liu et al., 2006; Rosa et al., 2015). As high-throughput studies have identified multiple candidate binding partners that have not yet been investigated (Boxem et al., 2008; Brajenovic et al., 2004; Giot et al., 2003; Grossmann et al., 2015; Hein et al., 2015; Huttlin et al., 2015; Lenfant et al., 2010; Luck et al., 2020; Waaijers et al., 2016), additional interactions important for the functioning of Par6 and for linking cortical polarity to other processes involved in epithelial polarization likely remain to be discovered.

Despite the conserved requirements for Par6–aPKC and Par3 in epithelial cells there are important context and cell-type dependent differences in the functioning of these polarity proteins (Pickett et al., 2019; St Johnston, 2018). For example, in Drosophila, Bazooka (Par3) is not required for junction positioning or polarization of cells in the follicular epithelium (Pickett et al., 2019; Shahab et al., 2015), and in the adult Drosophila midgut, the canonical Par, Crumbs, and Scribble polarity modules are all dispensable for apical–basal polarity (Chen et al., 2018). In *C. elegans*, requirements for PAR-3 and PAR-6 in embryonic epithelia also vary. PAR-6 appears to be required for apical junction formation in all epithelia, including the epidermis, intestine, foregut, and pharyngeal arcade cells (Montoyo-Rosario et al., 2020; Totong et al., 2007; Von Stetina and Mango, 2015; Von Stetina et al., 2017). However, while arcade cells show a complete lack of polarization upon PAR-6 loss, foregut, intestinal, and epidermal epithelial cells still establish an apical domain (Totong et al., 2007; Von Stetina and Mango, 2015). PAR-3 is required for apical junction formation in embryonic intestinal and pharyngeal epithelia, but not in epidermal epithelial cells (Achilleos et al., 2010).

Studies of PAR-6, PKC-3, and PAR-3 in *C. elegans* have largely focused on embryonic tissues. Here, we make use of targeted protein degradation to investigate the role of PAR-6, PKC-3, and PAR-3 in larval epithelia of *C. elegans*. Ubiquitous depletion of PAR-6 and PKC-3, but not PAR-3, resulted in a larval growth arrest, demonstrating that these proteins are required for larval development. Through tissue-specific depletion, we identified an essential role for PAR-6 and PKC-3 in the *C. elegans* epidermis. Depletion in this tissue caused growth arrest, a cessation of the molting cycle, and severe defects in the division pattern of the epidermal seam cells. We also observed defects in the maintenance of apical cell junctions, and a failure to exclude LGL-1 from the apical domain. Finally, we identified a novel role for PAR-6 in organizing non-centrosomal microtubule arrays in the epidermis. Epidermal depletion of PAR-6 led to defects in the localization of the microtubule organizer NOCA-1/Ninein, as well as of the γ-tubulin ring complex component GIP-1, and of the sole Patronin/CAMSAP/Nezha homolog PTRN-1. Microtubule defects in a *noca-1* mutant closely resembled those in PAR-6 depleted animals, including the loss of GIP-1 localization. As NOCA-1 physically interacts with PAR-6, we conclude that PAR-6 likely organizes non-centrosomal microtubule arrays through localization of NOCA-1.

## Results

### PAR-6 and PKC-3 are essential for larval development

To investigate the role of PAR-6, PKC-3, and PAR-3 in larval development, we made use of the auxin-inducible degradation (AID) system. The AID system enables targeted degradation of AID-degron tagged proteins through expression of the plant-derived auxin-dependent E3 ubiquitin ligase specificity factor TIR1 (Zhang et al., 2015) (Fig. 1A). Using CRISPR/Cas9, we inserted sequences encoding the AID-degron and the green fluorescent protein (GFP) into the endogenous *par-6, pkc-3*, and *par-3* loci, such that all known isoforms of each protein are tagged (Fig. 1B). PAR-6 was tagged at the shared C-terminus, and PKC-3 at the N-terminus. The *par-3* locus encodes two groups of splice variants that use two alternative start sites; hence we inserted the GFP–AID tag at both start sites. To examine if the presence of the GFP–AID tags interfered with protein function, we examined the growth rates of the tagged strains. Homozygous animals were viable and showed the same growth rates as wild-type, indicating that the proteins are still functional (Fig. 1C–E). We also observed localization of each protein at the apical membrane domain of epithelial tissues, including the pharynx, excretory canal, intestine and epidermis (Fig. 1F, G). This localization pattern matches previous observations in *C. elegans* larvae (Li et al., 2010a; Li et al., 2010b), and further indicates that the GFP–AID tag does not interfere with protein functioning.

**Fig. 1.**
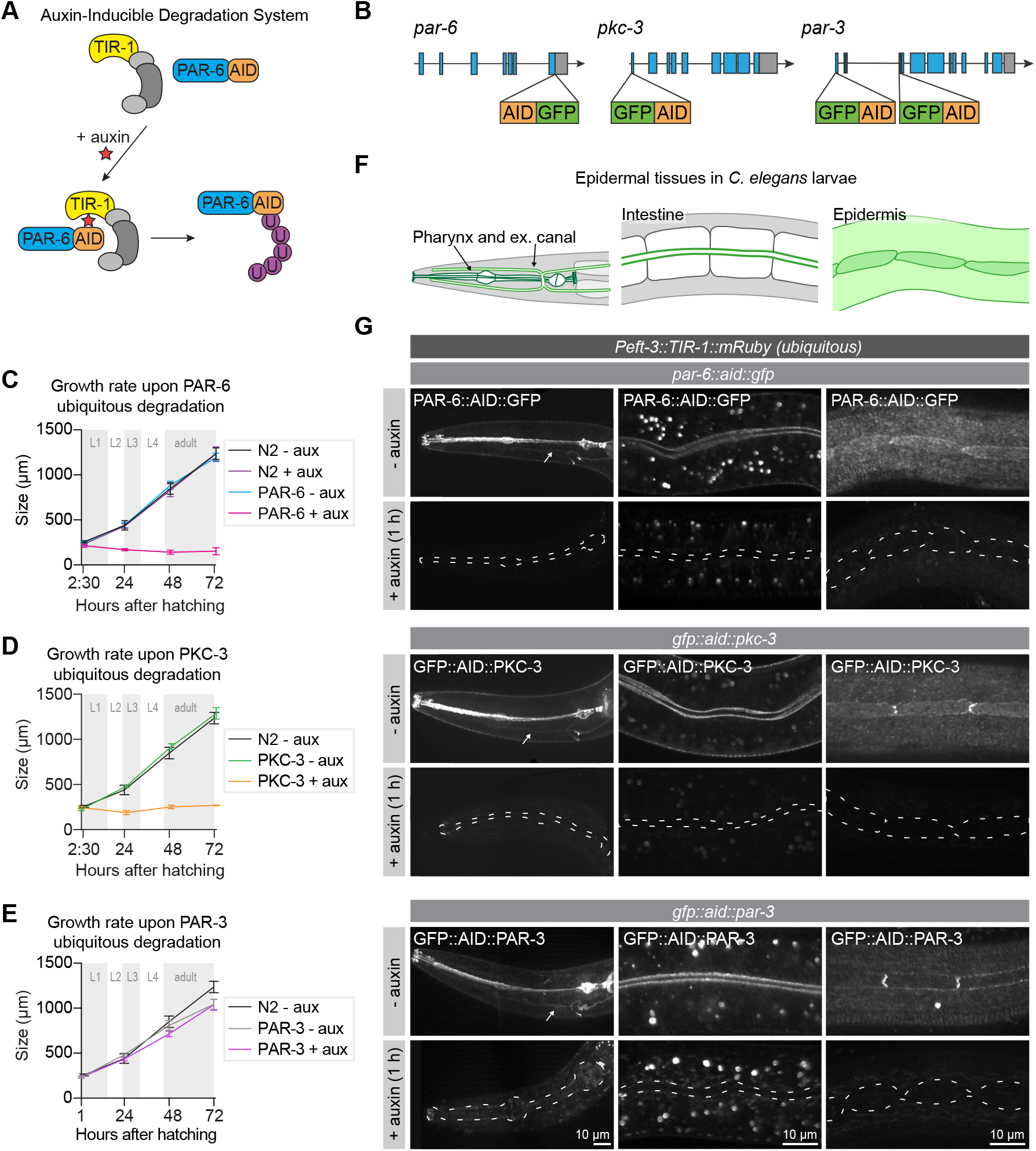
PAR-6 and PKC-3 are essential for larval development. **(A)** Overview of the AID system, which enables targeted degradation of AID-tagged proteins by the plant-derived E3 ubiquitin ligase specificity factor TIR1 upon addition of auxin. **(B)** Schematic representation of endogenous tagging of *par-6, pkc-3*, and *par-3* loci with sequences encoding a green fluorescent protein (GFP) and auxin-inducible degradation degron (AID) tag. **(C–E)** Growth curves of N2, *par-6::aid::gfp, gfp::aid::pkc-3*, and *gfp::aid::par-3* animals in absence (− aux) or presence (+ aux) of 4 mM auxin. Data show mean ± SD. *n* = 6, 7, 8, and 8 for N2 - aux; 6, 7, 9, and 9 for N2 + aux; 7, 6, 9, and 9 for PAR-6 - aux; 8, 6, 7, and 9 for PAR-6 + aux; 22, 11, 10, and 14 for PKC-3 - aux; 19, 14, 9, and 10 for PKC-3 + aux; 10, 10, 10, and 10 for PAR-3 - aux, and 10, 10, 10, and 10 for PAR-3 + aux. **(F)** Graphical representation of larval epithelial tissues in *C. elegans*. Green indicates localization of PAR-6, PKC-3, and PAR-3. **(G)** Distribution of GFP::AID-tagged PAR-6, PKC-3, and PAR-3 in different larval tissues in absence (− auxin) or presence (+ auxin) of 4 mM auxin. Images are maximum projections. Dashed lines in - auxin panels outline pharynx (left panel), intestinal lumen (middle panel) or seam cells (right panel). White arrows point to the excretory canals.

To investigate the role of PAR-3, PAR-6 and PKC-3 in larval development we degraded each protein using a ubiquitously expressed TIR1 under the control of the *eft-3* promoter (Zhang et al., 2015). We tested the efficiency of protein degradation by exposing synchronized L3 larvae to auxin and examining protein expression. Apical enrichment of PAR-3, PAR-6, and PKC-3 became indistinguishable from background fluorescence within one hour of exposure to 4 mM auxin in the pharynx, excretory canal, intestine, and epidermis (Fig. 1G). To examine if the depletion of PAR-6, PKC-3, or PAR-3 affected larval development, we degraded each protein by addition of auxin at hatching and measured animal growth rates. Ubiquitous degradation of PAR-3 did not cause a defect in larval growth, and animals developed into morphologically normal looking and fertile adults (Fig. 1E). This lack of a visible phenotype may indicate that the functions of PAR-3 are not essential for larval development. Alternatively, despite visual absence of GFP::AID::PAR-3, degradation may be incomplete, or animals may express unpredicted non-tagged protein isoforms. In contrast to PAR-3, depletion of PAR-6 or PKC-3 caused a striking growth arrest with animals not developing beyond L1 size (Fig. 1C, D). Thus, PAR-6 and PKC-3 are essential for early larval development, and we focused our further analysis on PAR-6 and PKC-3.

### PAR-6 and PKC-3 are essential in the larval epidermis, but not in the intestine

We next wanted to determine which larval tissue or tissues are severely affected by the loss of PAR-6/ PKC-3 and contribute to the growth arrest. We focused on the two major epithelial organs: the intestine and the epidermis. The intestine is an epithelial tube formed in embryogenesis by 20 cells, which do not divide during larval development. PAR-6 and PKC-3 are highly enriched at the apical luminal domain (Fig. 2A). The epidermis consists of two cell types: hypodermal cells and seam cells.

**Fig. 2.**
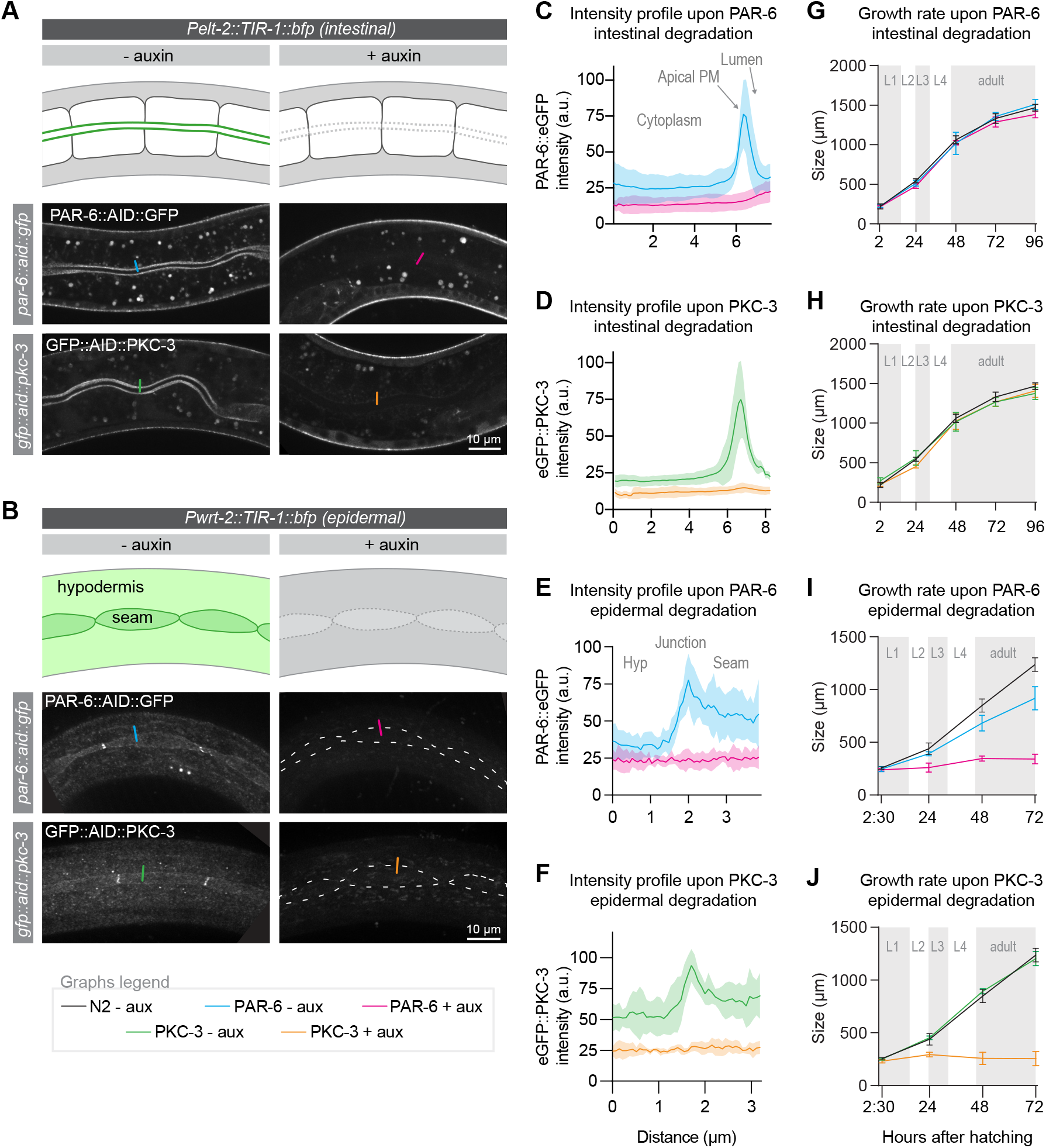
PAR-6 and PKC-3 are essential in the epidermis to support larval growth. **(A, B)** Distribution of PAR-6::AID::GFP and GFP::AID::PKC-3 in the intestine (A) and epidermis (B) in absence (− auxin) or presence (+ auxin) of 1 mM auxin. Images are maximum projections of the luminal domain for the intestine, and the apical domain for the epidermis. Drawings are schematic representation of the area imaged, with the localization of PAR-6 and PKC-3 indicated in green shades. Greys indicate absence of PAR-6 and PKC-3. **(C–F)** Quantification of apical GFP fluorescence intensity at the intestinal lumen and the hyp7–seam cell junction (indicated by colored lines in A, B) in *par-6::aid::gfp* and *gfp::aid::pkc-3* animals in the absence (− aux) or presence (+ aux) of 1 mM auxin. Solid lines and shading represent mean ± SD. For the intestine, n = 10 animals for PAR-6 - aux, PAR-6 + aux, PKC-3 - aux, and PKC-3 + aux. For the epidermis, n = 8 animals for PAR-6 - aux, 6 for PAR-6 + aux, 5 for PKC-3 + aux, and 5 for PKC-3 - aux. **(G–J)** Growth curves of N2, *par-6::aid::gfp*, and *gfp::aid::pkc-3* animals in absence (− aux) or presence (+ aux) of 4 mM auxin. Solid lines and shading represent mean ± SD. In G and H, degradation was induced in the intestine, and in I and J in the epidermis. In the intestine, n = 13, 10, 13, 14, and 12 for N2 - aux; 7, 7, 7, 5, and 9 for PAR-6 - aux; 6, 6, 6, 5, and 7 for PAR-6 + aux; 8, 7, 8, 4, and 9 for PKC-3 - aux; and 8, 7, 8, 8, and 8 for PKC-3 + aux. In the epidermis, *n* = 6, 7, 8, and 8 for N2 - aux; 6, 5, 11, and 8 for PAR-6 - aux; 5, 10, 8, and 9 for PAR-6 + aux; 7, 7, 10, and 8 for PKC-3 - aux; and 8, 7, 12, and 13 for PKC-3 + aux.

The hypodermal syncytial cell hyp7 covers most of the body. Embedded within hyp7 are two lateral rows of epithelial seam cells, which contribute multiple nuclei to hyp7 through asymmetric divisions in each larval stage. PAR-6 and PKC-3 localize to the apical domain of the seam cells and hyp7, and are enriched at the seam/seam and seam/hyp7 junctions (Fig. 2B).

To enable tissue-specific depletion of PAR-6 and PKC-3, we generated single-copy integrant lines expressing TIR1 in the intestine and epidermal lineages, using the tissue-specific promoters *Pelt-2* and *Pwrt-2*, respectively. In both tissues, protein depletion occurred within one hour of addition of 1 mM auxin (Fig. 2A–F). To determine the contribution of the intestine and epidermis to the larval growth defects we observed above, we measured the growth rate of animals depleted of PAR-6 or PKC-3 in each tissue. Depletion of either protein from the intestine did not result in a growth delay or in obvious defects in morphology of the intestine (Fig. 2G, H). Simultaneous depletion of PAR-6 and PKC-3 also did not result in a growth delay or visible abnormalities in the intestine (Fig. S1A). These data indicate that PAR-6 and PKC-3 are not essential for the functioning and homeostasis of the larval intestine, though we cannot exclude that very low protein levels that we were not able to detect by fluorescence microscopy are sufficient in this tissue.

A recent study in Drosophila indicated a potential redundancy between aPKC and the kinase Pak1 (Aguilar-Aragon et al., 2018). To determine if a similar redundancy was the cause of the absence of intestinal PKC-3 depletion phenotypes, we combined depletion of PKC-3 with inactivation of *C. elegans pak-1* by RNAi. However, we again did not observe a delay in larval growth (Fig. S1B).

In contrast to the intestine, depletion of PAR-6 or PKC-3 from hatching in the epidermis caused an early larval growth arrest, as observed with ubiquitous degradation (Fig. 2I, J). Thus, PAR-6 and PKC-3 play an essential role in the functioning and/or development of epidermal larval epithelia. We did notice that animals with ubiquitous PAR-6 or PKC-3 depletion appeared more sick than epidermal depleted animals, indicating that the functions of PAR-6 and PKC-3 are not limited to the epidermis.

### Cell autonomous and non-autonomous roles for PAR-6 and PKC-3 in molting, seam cell divisions and seam cell morphology

One of the major functions of the epidermal epithelium is the synthesis and apical secretion of cuticle components, and defects in this tissue can lead to molting defects (Chisholm and Xu, 2012; Lažetić and Fay, 2017). As molting defects can be accompanied by a growth arrest (Brooks et al., 2003; Lažetić and Fay, 2017; Russel et al., 2011; Yochem et al., 1999), we explored the possibility that PAR-6 depleted animals are molting defective. By Nomarski differential interference contrast (DIC) microscopy, we observed incompletely released cuticles 30 hours past exposure to auxin, indicative of molting defects (Fig. 3A). To examine molting progression in more detail, we used a transcriptional reporter expressing GFP from the *mlt-10* promoter (Meli et al., 2010). *mlt-10* expression cycles, increasing during molting and decreasing during the inter-molt. Upon epidermal degradation of PAR-6 from hatching, *mlt-10* driven GFP levels did not cycle and remained low (Fig. 3B, C), indicating a perturbation of the molting cycle. Together, these results demonstrate that PAR-6 performs essential functions in the epidermis required to support remodeling of the *C. elegans* cuticle.

**Fig. 3.**
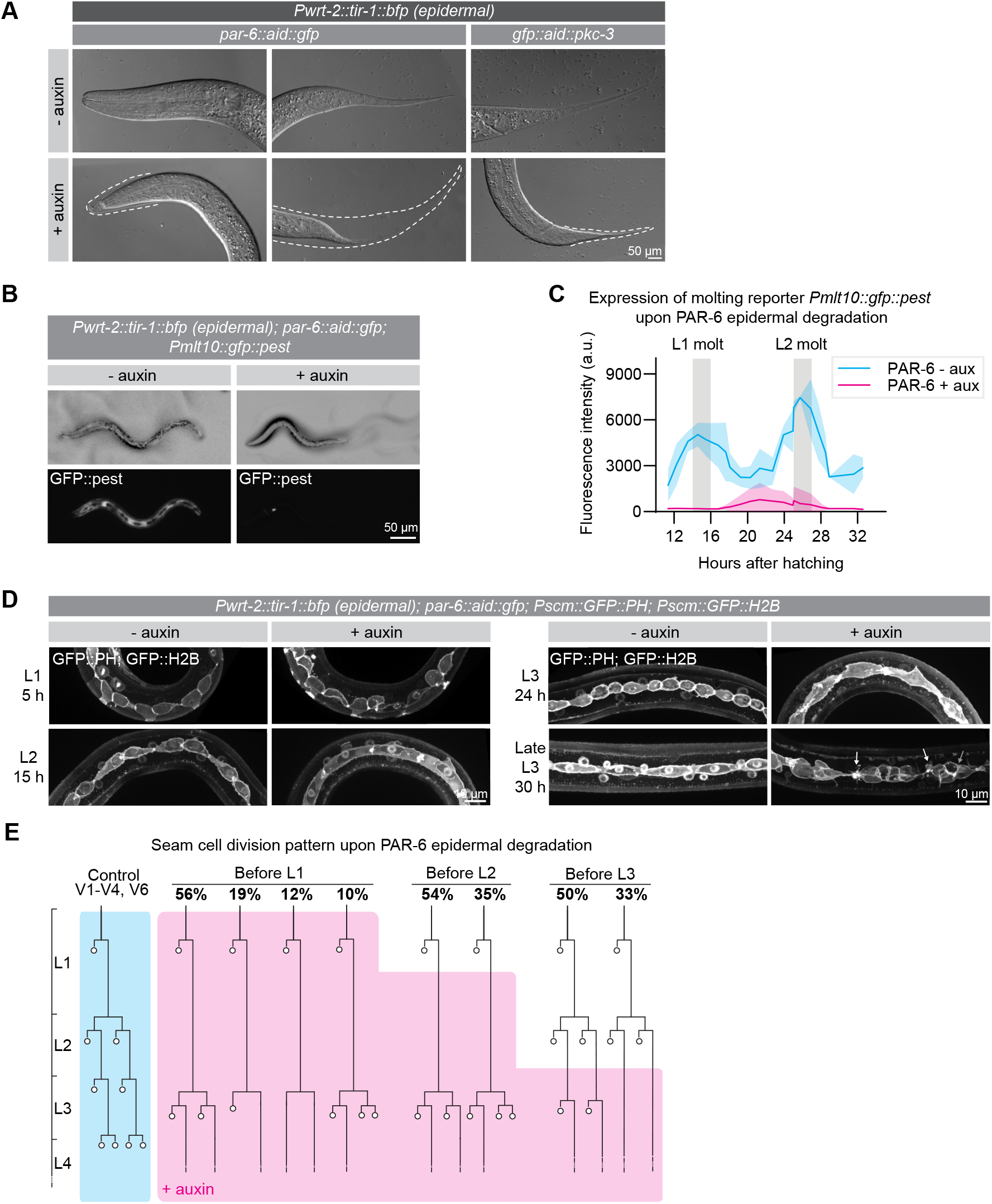
PAR-6 and PKC-3 are required in the epidermis for molting and seam cell development. **(A)** DIC microscopy images of molting defects upon epidermal depletion of PAR-6 or PKC-3. Animals were grown in absence (−auxin) or presence (+auxin) of 1 mM auxin since hatching, and images were taken 30 h after hatching. Dotted lines outline detached but unreleased cuticle in the pharynx and in the tail. Defects are observed in ~50% of the animals. **(B)** Expression of the molting reporter *Pmlt-10::gfp::pest* in *par-6::aid::gfp* animals in the absence (− auxin) or presence (+ auxin) of 1 mM auxin at 22 h of post-embryonic development. **(C)** Quantification of *Pmlt-10::gfp::pest* expression from 11 h to 32 h of post-embryonic development (mean fluorescence intensity ± SD) in *par-6::aid::gfp* animals in absence (− aux) or presence from hatching (+ aux) of 1 mM auxin. Measurements were done every hour. Each data point is an average of 3–12 measurements, with an average of 8 measurements per data point. **(D)** Examples of seam cell division and morphology defects observed upon depletion of PAR-6::GFP::AID. Seam cells are visualized by expression of nuclear H2B::GFP and membrane-bound PH::GFP markers (Wildwater et al., 2011). Arrows indicate membrane protrusions and arrowhead indicates abnormal division plane orientation. **(E)** Seam cell division pattern in *par-6::aid::gfp* animals in absence (control, blue) or presence (+ auxin, magenta) of 1 mM auxin. For the control, *n* = 14, 75, 40, and 28 animals for the L1, L2, L3 and L4 divisions. For before L1, n = 17 animals for the L1 and 143 animals for the delayed L2 divisions. For before L2, n = 91 animals. For before L3, n = 40 animals.cell junction, normalized to the background intensity of each animal measured in the hypodermis. n = 6 animals for both conditions. **(E, F)** Junction organization visualized by DLG-1::mCherry expression in *par-6::aid::gfp* or *gfp::aid::pkc-3* animals in the absence (− auxin) or presence (+ auxin) of 1 mM auxin for 6 (E) or 24 (F) hours. **(G)** Graphical representation of junctional defects in the seam cells upon PAR-6 or PKC-3 degradation.15 for PAR-6 deg, 14 for PKC-3 deg and 13 for *noca-1(ok3692).* Bars show mean ± SD **(E)** Microtubule growth visualized by maximum intensity projection of the plus end marker EBP-2::GFP in absence (− auxin) or presence (+ auxin) of 1 mM auxin for 1 h. To match the age of animals in (C), we depleted PAR-6 for 1 h starting with 23 h old L2 animals. **(F)** EBP-2 comet density. n = 12 animals for control and PAR-6 deg, 8 for *noca-1(ok3692)*, and 8 for PAR-6 deg + rescue. Bars show mean ± SD **(G)** Microtubule growth rate. n > 400 comets. Bars = mean ± SD **(H)** Quantification of microtubule growth orientation. Vertical axis: left/right orientation; horizontal axis: anterior/posterior orientation. n = 150 comets. Bars = mean ± SD. Tests of significance: Tukey’s multiple comparisons test for D, and Dunn’s multiple comparisons test for F and G. ns = not significant, * = P ≤ 0.05, ** = P ≤ 0.01, *** = P ≤ 0.001, **** = P ≤ 0.0001.

We next examined the effects of PAR-6 depletion on the stereotypical division pattern of the seam cells. In every larval stage, an asymmetric cell division creates a new seam cell daughter and a cell that differentiates to form neurons or fuse with hyp7 (Chisholm and Hsiao, 2012) (Fig. 3E, blue shaded lineage tree). In the second larval stage, a symmetric division precedes the asymmetric division to double the number of seam cells. Depletion of PAR-6 directly after hatching did not disrupt the L1 asymmetric division, but the divisions that normally take place in the L2 stage were severely delayed (Fig. 3D, E). At the time when control animals were already undergoing the L3 divisions, L2-stage divisions had still not taken place in PAR-6 depleted animals. Eventually, a next round of divisions did take place, but we observed various deviations from the normal L2 division pattern, including division failures and abnormal differentiation and fusion with hyp7. We did not observe any L3 divisions (Fig. 3E). We also observed numerous morphological abnormalities such as membrane protrusions, blebs, and abnormal division plane orientation (Fig. 3D). Exposure of synchronized populations to auxin starting after the L1 or L2 divisions resulted in similar defects, indicating that the seam cell lineage requires the functioning of PAR-6 throughout development (Fig. 3E).

Expression of TIR1 under the *wrt-2* promoter results in degradation of target proteins in both hyp7 and the seam cells. While both tissues contribute to cuticle production, severe molting defects can result from the loss of hyp7-specific proteins, such as the NIMA-related kinases NEKL-2,3 (Yochem et al., 2015). It is possible therefore the seam cell defects are a secondary consequence of the molting defect. To address this, we expressed an exogenous copy of *par-6:mCherry* lacking the degron sequence in hyp7 using the hypodermis-specific *dpy-7* promoter (Gilleard et al., 1997). In combination with auxin-induced depletion of PAR-6::GFP::AID by *wrt-2* driven TIR-1, this results in absence of PAR-6 only from the seam cells. Hypodermal specific expression of *par-6::mCherry* rescued the cuticle defects and seam cell division delay observed upon PAR-6 epidermal degradation, and partially rescued the growth arrest (Fig. S2A–D). However, seam cell morphology defects and the abnormal cell division plane were not restored (Fig. S2C). Thus, PAR-6 expression in the hypodermis is sufficient to support molting and larval growth. Furthermore, at least some of the seam cell defects we observed, most notably the timing defect, appear to be a secondary consequence of hypodermal or molting defects. The fact that the growth arrest and seam abnormalities were not fully rescued may indicate cell autonomous roles for PAR-6 in the seam, or alternatively that the *Pdpy-7::par-6::mCherry* rescue construct is not fully functional.

To determine the effects of a molting arrest on seam cell divisions via an alternative approach, we used a CRISPR-tagged NEKL-2::AID strain that arrests molting upon auxin addition (Joseph et al., 2020). NEKL-2 depletion caused defects in the morphology of seam cells, as well as reduced cell division, confirming that abnormalities in the hypodermis can affect the seam cells (Fig. S2E, F). However, in contrast to PAR-6 depletion, not all seam cell divisions were affected, indicating that a block in molting does not fully explain the seam cell division delay in PAR-6 depleted animals.

Taken together, our data shows that PAR-6 and PKC-3 have essential roles in the *C. elegans* epidermis to support cuticle homeostasis and molting. Restoring *par-6(+)* expression specifically in the hypodermis revealed the importance of PAR-6 for this cell type and indicate that the hypodermis and/or molting are essential for proper seam cell development. Finally, PAR-6 may perform cell autonomous functions in the seam cells that contribute to proper seam cell shape and cell division plane.

### PAR-6 and PKC-3 mediate apical LGL-1 exclusion and promote junction integrity in the larval epidermis

As one of the major functions of the apical PAR complex is to mediate the exclusion of basolateral proteins from the apical domain, we next examined the effects of PKC-3 depletion on two key aPKC target genes: LGL-1/Lgl and PAR-1. Both proteins are direct aPKC targets in epithelia, and in the *C. elegans* zygote their anterior exclusion is mediated by PKC-3 (Beatty et al., 2010; Betschinger et al., 2003; Doerflinger et al., 2010; Hoege et al., 2010; Hurov et al., 2004; Motegi et al., 2011; Plant et al., 2003; Ramanujam et al., 2018; Yamanaka et al., 2003). For these experiments we made use of integrated LGL-1::GFP transgene (Waaijers et al., 2015) and an endogenously tagged PAR-1::GFP fusion.

Depletion of PKC-3 in the intestine did not result in apical invasion of LGL-1 (Fig. S3A). In contrast, degradation of PKC-3 in the epidermis resulted in clear apical LGL-1 localization in the seam cells within 6 hours of auxin addition (Fig. 4A, B). Degradation of PKC-3 in the epidermis did not result in apical PAR-1 localization (Fig. 4C, D). Instead, prolonged depletion of PKC-3 for 24h resulted in fragmentation of the normally contiguous PAR-1 signal at cell junctions, which may reflect an indirect effect of PKC-3 on junction organization (Fig. 4C). These results demonstrate that PKC-3 is necessary to maintain the basolateral localization of LGL-1 in the seam cells, but not the intestine. In contrast, the apical exclusion of PAR-1 is not solely mediated by aPKC, though the organization of PAR-1 at cell junctions does depend on PKC-3.

**Fig. 4.**
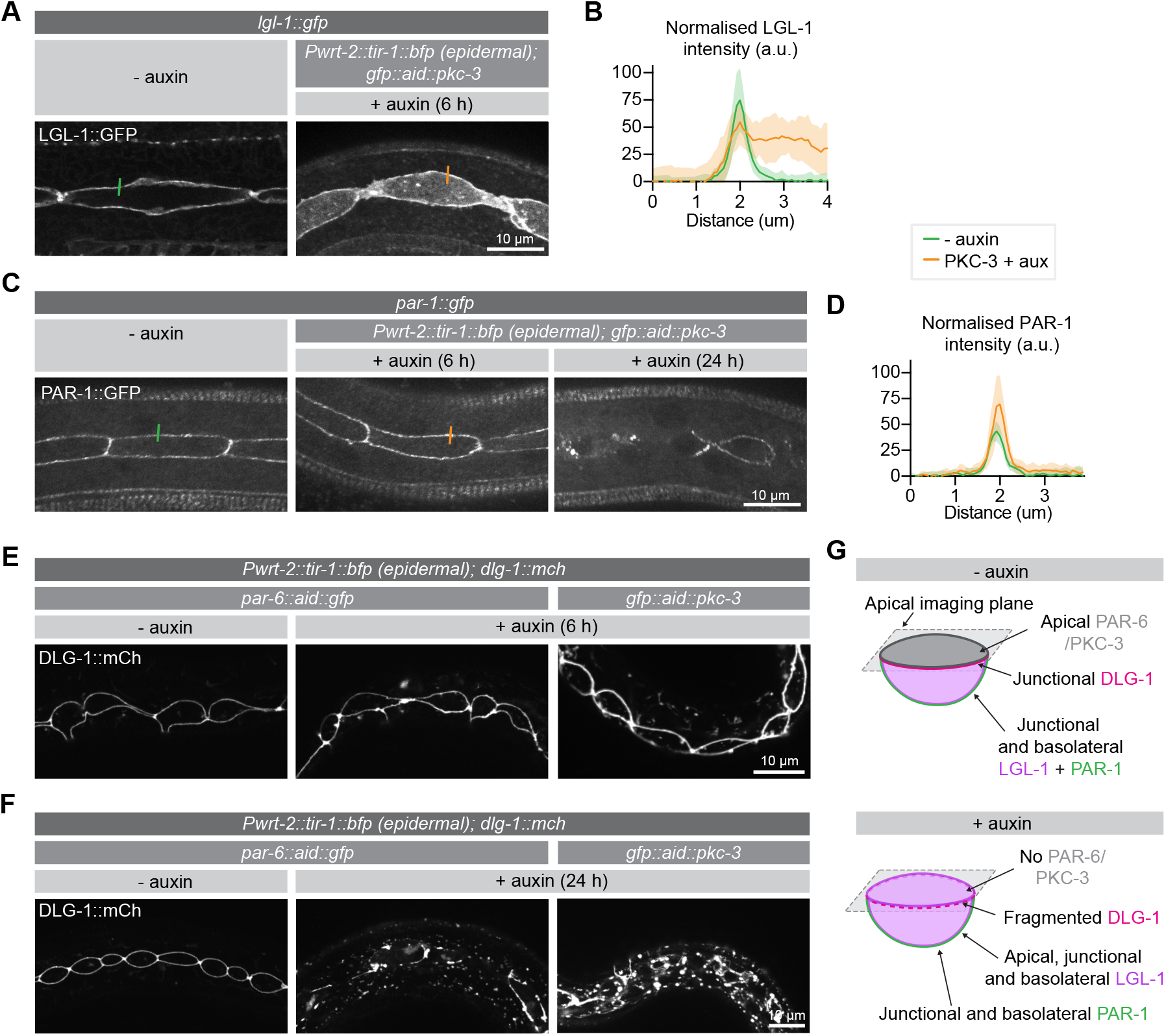
PKC-3 excludes LGL-1 from the apical cortex and, together with PAR-6, regulates junctions. **(A, B)** Distribution and quantification of LGL-1::GFP in the epidermis of *lgl-1::gfp* animals without auxin and in *lgl-1::gfp; gfp::aid::pkc-3; Pwrt-2::tir-1::bfp* animals in the presence of 4mM auxin. Images are maximum projections of the apical domain. Quantifications shows mean apical GFP fluorescence intensity ± SD at the hyp7–seam cell junction, normalized to background intensity of each animal measured in the hypodermis. n = 7 animals for both conditions. **(C, D)** Distribution and quantification of PAR-1::GFP in the epidermis in *par-1::gfp* animals without auxin and in *par-1::gfp; gfp::aid::pkc-3; Pwrt-2::tir-1::bfp* animals in the presence of 4mM auxin. Images are maximum projections of the apical domain. Quantifications show mean apical GFP fluorescence intensity ± SD at the hyp7–seam cell junction, normalized to the background intensity of each animal measured in the hypodermis. n = 6 animals for both conditions. **(E, F)** Junction organization visualized by DLG-1::mCherry expression in *par-6::aid::gfp* or *gfp::aid::pkc-3* animals in the absence (− auxin) or presence (+ auxin) of 1 mM auxin for 6 (E) or 24 (F) hours. **(G)** Graphical representation of junctional defects in the seam cells upon PAR-6 or PKC-3 degradation.

In embryonic epithelia, PAR-6 and PKC-3 are essential for proper junction formation, with loss of either protein resulting in fragmented cell junctions (Montoyo-Rosario et al., 2020; Totong et al., 2007). To investigate the requirement for PAR-6 and PKC-3 in junction integrity in larval epithelia, we assessed the localization of an endogenous GFP fusion of the junctional protein DLG-1 upon degradation of PAR-6 or PKC-3 from hatching. In control animals not exposed to auxin, DLG-1 displays the typical ladder-like intestinal junction pattern and forms a continuous apical belt around the seam cells (Figs S3B and 4E, F). Upon degradation of PAR-6 in the intestine, we did not observe junctional defects (Fig. S3B). We also did not observe any changes to the DLG-1 localization pattern in the epidermis after 6 hours of PAR-6 or PKC-3 depletion (Fig. 4E). However, after 24 h of degradation, DLG-1 no longer localized in a uniform band around the seam cells but appeared fragmented, with aggregates of bright DLG-1 interspersed with areas lacking fluorescent signal (Fig. 4F). We also observed fluorescent aggregates in the hypodermis (Fig. 4F). Thus, as in the embryo, PAR-6 and PKC-3 are essential for junction integrity in the epidermis. The fact that junctional defects took 24 h to develop, compared to 6 h for LGL-1 mislocalization, points to an inherent stability of cell junctions.

Finally, we investigated the localization dependencies between PAR-6, PKC-3 and PAR-3. Several studies demonstrated that PAR-6 and PKC-3 co-localize throughout development, and are mutually dependent on each other for their asymmetric localization (Bossinger et al., 2001; Leung et al., 1999; McMahon et al., 2001; Nance and Priess, 2002; Nance et al., 2003; Tabuse et al., 1998; Totong et al., 2007). Moreover, binding of PAR-6 to PKC-3 is required for apical localization of PAR-6, including in larval epithelia (Li et al., 2010a). Not surprisingly therefore, degradation of PAR-6 resulted in the loss of PKC-3 from the apical domain of the seam cells, and degradation of PKC-3 similarly disrupted PAR-6 localization (Fig. S4A, B). These disruptions occurred rapidly, within 1 h of auxin addition. Depletion of PAR-6 in the intestine also caused rapid loss of PKC-3 from the apical plasma membrane, demonstrating that, despite the lack of intestinal phenotypes, PAR-6 and PKC-3 do form an apical protein complex (Fig. S3C, D). When we followed the apical loss of PKC-3 in the intestine over time, we observed similar dynamics of PAR-6 depletion and PKC-3 loss (Video S1). Our results thus confirm the interdependency between PAR-6 and PKC-3. Surprisingly, the effects of PAR-6 and PKC-3 depletion on PAR-3 were distinct: the apical localization of PAR-3 in the seam cells depended on the presence of PKC-3, but not PAR-6 (Fig. S4C, D). Finally, we examined the effects of PAR-3 degradation on PAR-6 and PKC-3 in the seam cells. We observed partially reduced levels of PAR-6, but no effect on PKC-3 (Fig. S4E, F). Thus, PAR-6 and PKC-3 may not form an obligatory dimer under all conditions.

### PAR-6 and PKC-3 control the organization of non-centrosomal microtubule arrays in the hypodermis

The loss of PAR-6 or PKC-3 affected several epidermal processes in which cytoskeletal elements play important roles, including molting, seam cell divisions, and maintaining proper seam cell morphology. The PAR proteins play essential roles in organizing the actomyosin cytoskeleton and microtubules in different settings, including asymmetric cell division, neuronal differentiation, and epithelial polarization (Goldstein and Macara, 2007; Rodriguez-Boulan and Macara, 2014; St Johnston, 2018). We therefore investigated if PAR-6 degradation affects the organization of actin or microtubules in the epidermis. To assess the organization of the actin cytoskeleton we used an epidermal transgene expressing the actin-binding-domain of *vab-10* fused to *mCherry* (Gally et al., 2009). We depleted PAR-6 from hatching and examined actin organization after 24 h in late L2 larvae. Consistent with previous observations (Costa et al., 1997), we observed prominent circumferential actin bundles in hyp7, strong enrichment of actin along the hyp7/seam junctions, and largely anterior/posteriorly organized actin within the seam cells of control animals at this stage (Fig. 5A). Upon PAR-6 depletion, actin organization appeared largely undisturbed in both the seam cells and hypodermis (Fig. 5A), and actin bundles in hyp7 remained perpendicular to the seam cells (Fig. 5B). These data indicate that PAR-6 does not play a major role in regulating the actin cytoskeleton in the larval epidermis.

**Fig. 5.**
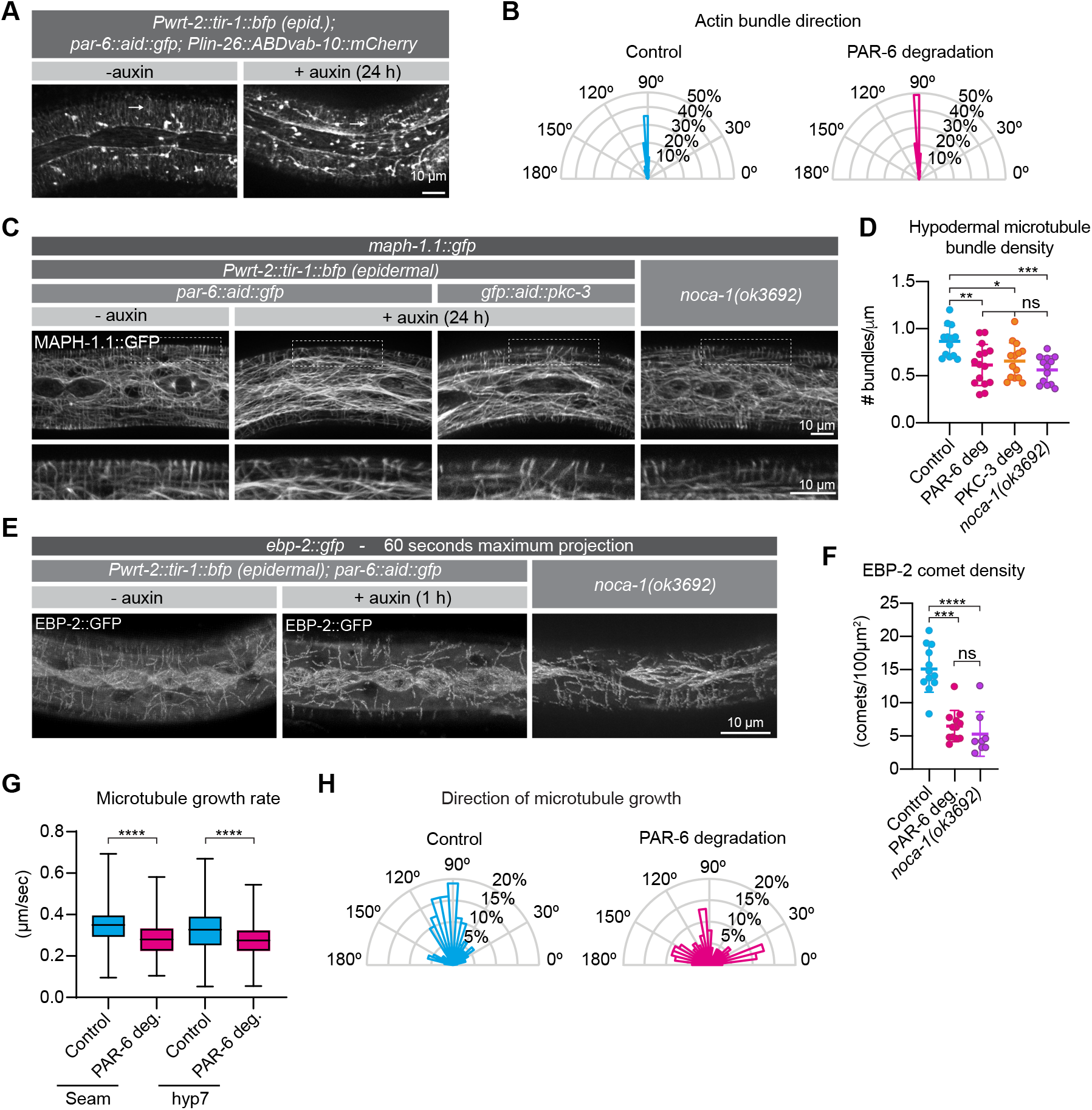
PAR-6 and PKC-3 control microtubule organization in the *C. elegans* epidermis. **(A)** Actin organization visualized by the *Plin-26::ABDvab-10::mCherry* reporter in *par-6::aid::gfp* animals in absence (− auxin) or presence (+ auxin) of 1 mM auxin for 24 hours. **(B)** Quantification of actin bundle orientation. Angle is measured relative to the anterior (180°) – posterior (0°) axis. n = 100 bundles in 5 animals per condition. **(C)** Microtubule organization of the indicated genotypes visualized by MAPH-1.1::GFP in absence (− auxin) or presence (+ auxin) of 1 mM auxin for 24 hours. Images are maximum intensity projections. **(D)** Hypodermal microtubule bundle density. n = 13 animals for control, 15 for PAR-6 deg, 14 for PKC-3 deg and 13 for *noca-1(ok3692).* Bars show mean ± SD **(E)** Microtubule growth visualized by maximum intensity projection of the plus end marker EBP-2::GFP in absence (− auxin) or presence (+ auxin) of 1 mM auxin for 1 h. To match the age of animals in (C), we depleted PAR-6 for 1 h starting with 23 h old L2 animals. **(F)** EBP-2 comet density. n = 12 animals for control and PAR-6 deg, 8 for *noca-1(ok3692)*, and 8 for PAR-6 deg + rescue. Bars show mean ± SD **(G)** Microtubule growth rate. n > 400 comets. Bars = mean ± SD **(H)** Quantification of microtubule growth orientation. Vertical axis: left/right orientation; horizontal axis: anterior/posterior orientation. n = 150 comets. Bars = mean ± SD. Tests of significance: Tukey’s multiple comparisons test for D, and Dunn’s multiple comparisons test for F and G. ns = not significant, * = P ≤ 0.05, ** = P ≤ 0.01, *** = P ≤ 0.001, **** = P ≤ 0.0001.

We next inspected the organization of the microtubule cytoskeleton using an endogenously GFP tagged variant of the microtubule-binding protein MAPH-1.1 (Waaijers et al., 2016). We degraded PAR-6 in the epidermis from hatching and assessed the organization of epidermal microtubule arrays after 24 h. In control animals, we observed highly ordered circumferential microtubule bundles in the dorsal and ventral sections of hyp7 underlying the muscle quadrants, and a microtubule meshwork in the lateral sections of hyp7 abutting the seam cells, as previously reported (Chuang et al., 2016; Costa et al., 1997; Taffoni et al., 2020; Wang et al., 2015) (Fig. 5C). In the seam cells the microtubule network was less well defined but also forms a meshwork (Fig. 5C). In PAR-6 depleted animals, we observed a significant reduction in the density of circumferential microtubule bundles in the hypodermis (Fig. 5C, D). Epidermal depletion of PKC-3 resulted in similar defects (Fig. 5C, D). To understand the cause of the reduced microtubule density, we investigated microtubule dynamics using an endogenous fusion of the microtubule plus-end tracking protein EBP-2^EB1^ to GFP. In control animals, EB1 comets moved along trajectories consistent with the organization of microtubule bundles in the epidermis, and both comet density and growth rates match previous reports (Fig. 5 E–G) (Chuang et al., 2016; Taffoni et al., 2020; Wang et al., 2015). Already within 1 h of inducing depletion of PAR-6, we observed reduced microtubule dynamics (Fig. 5E–G). The density of growing MTs was reduced by 56% (Fig. 5F), and microtubule growth rate was reduced by 14% in hyp7 and by 16% in the seam cells (Fig. 5G). These results suggest that the reduced density of microtubule bundles upon depletion of PAR-6 is the result of reduced growth or nucleation of microtubules. We also observed a defect in the directionality of microtubule growth. While 54% of the comets in control animals travel perpendicular to the seam cells (70–110°), this number is reduced to 24% upon PAR-6 degradation (Fig. 5H), consistent with the defects in organization observed with GFP::MAPH-1.1.

### PAR-6 controls microtubule organization through its interaction partner NOCA-1/Ninein and the γ-tubulin ring complex

Two large-scale protein–protein interaction mapping studies in *C. elegans* had identified the microtubule organizing protein NOCA-1 as an interaction partner of PAR-6 (Boxem et al., 2008; Lenfant et al., 2010). Affinity purification experiments showed that PAR-6 interacts with NOCA-1 through its PDZ domain (Lenfant et al., 2010), and we were able to confirm the PAR-6 PDZ interaction with NOCA-1 by yeast two-hybrid (Fig. S5). NOCA-1 functions together with γ-tubulin to assemble non-centrosomal microtubule arrays in multiple tissues, including the epidermis, and is thought to be a functional homolog of the vertebrate microtubule organizer Ninein (Green et al., 2011; Wang et al., 2015). NOCA-1 localizes to the apical cortex in seam cells, similar to the localization of PAR-6 (Figs 1C, 6A), but the mechanisms that mediate apical localization of NOCA-1 are currently not known. The physical interaction between PAR-6 and NOCA-1 prompted us to investigate if PAR-6 regulates non-centrosomal microtubule arrays through NOCA-1.

**Fig. 6.**
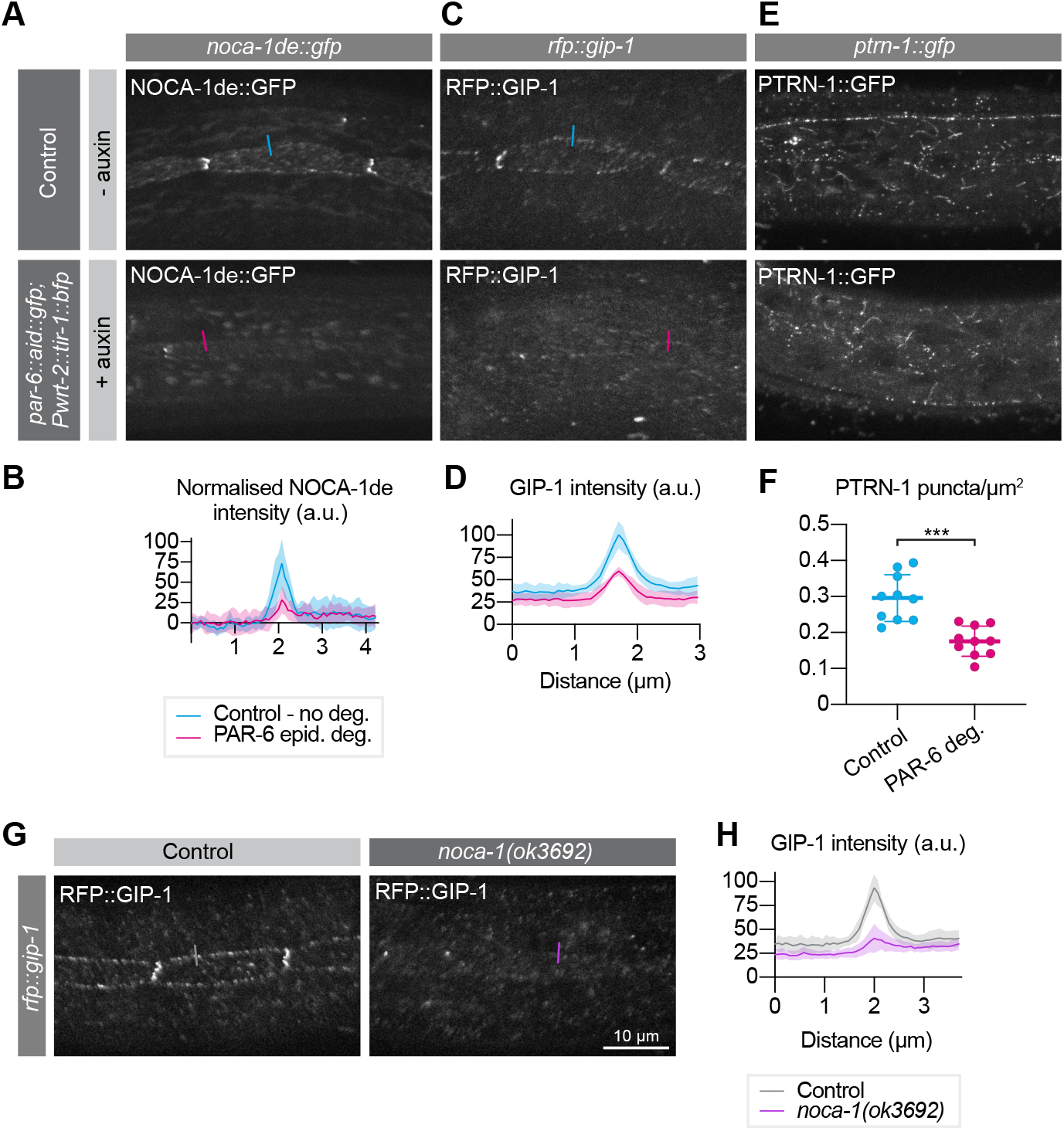
PAR-6 promotes the localization of its binding partner NOCA-1, as well as of GIP-1 and PTRN-1. **(A, B)** Distribution and quantification of NOCA-1de::GFP in the epidermis of *noca-1de::gfp* animals without auxin, and *noca-1de::gfp; par-6::aid::gfp; Pwrt-2::tir-1::bfp* animals in the presence of 4mM auxin. n = 9 animals for Control, and 10 for PAR-6 epid. deg. **(C, D)** Distribution and quantification of GIP-1::RFP in the epidermis of *gip-1::rfp* animals without auxin, and *gip-1::rfp; par-6::aid::gfp; Pwrt-2::tir-1::bfp* animals in the presence of 4mM auxin. n = 6 for Control and 6 for PAR-6 epid. deg. **(E, F)** Distribution and quantification of PTRN-1::GFP in the epidermis of *ptrn-1::gfp* animals without auxin, and *ptrn-1::gfp; par-6::aid::gfp; Pwrt-2::tir-1::bfp* animals in the presence of 4mM auxin. n = 10 for Control and 10 for PAR-6 deg. **(G, I)** Distribution and quantification of GIP-1::RFP in the epidermis of *gip-1::rfp* animals and *gip-1::rfp; noca-1(ok3692)*. n = 6 for Control and 6 for *noca-1(ok3692).* All images are maximum projections of the apical domain. Quantifications in B, D and H show mean apical GFP fluorescence intensity ± SD at the hyp7-seam cell junction (indicated by colored lines), normalized to background intensity of each animal measured in the hypodermis. Quantification in F shows mean PTRN-1::GFP puncta density ± SD. Tests of significance: unpaired t-test for F. *** = P ≤ 0.001.

We first examined the effect of epidermal PAR-6 depletion on the localization of NOCA-1. To visualize NOCA-1 we made use of an existing transgenic line that expresses the epidermis specific NOCA-1d and e isoforms fused to GFP from their endogenous promoter (Wang et al., 2015). In untreated control animals, we observed punctate localization of NOCA-1 in the epidermis, mostly clustered at the seam/seam and seam/hyp7 junctions, as previously observed (Fig. 6A) (Wang et al., 2015). Addition of auxin to induce epidermal PAR-6 degradation led to a 61% reduction in junctional levels of NOCA-1 within 6 hours (Fig. 6A, B), demonstrating that PAR-6 promotes the apical localization of NOCA-1.

NOCA-1 was reported to work together with γ-tubulin and redundantly with Patronin/PTRN-1 in controlling circumferential microtubule bundle organization in the hypodermis (Wang et al., 2015). We therefore examined the effect of PAR-6 depletion on the localization of PTRN-1 and GIP-1, a core component of the γ-tubulin ring complex (γ-TuRC) required to localize other γ-TuRC components to the apical non-centrosomal microtubule organizing center (ncMTOC) in the embryonic intestine (Sallee et al., 2018). To visualize PTRN-1 and GIP-1 we used endogenous PTRN-1::GFP and RFP::GIP-1 fusion proteins. GIP-1 localized in a punctate pattern at the seam/seam and seam/hyp7 junctions, similar to NOCA-1 (Fig. 6C) (Sallee et al., 2018; Wang et al., 2015). PTRN-1 also localized in a punctate pattern, but dispersed through the epidermis and lacking the junctional enrichment seen for NOCA-1 and GIP-1 (Fig. 6E) (Wang et al., 2015). Upon PAR-6 degradation, junctional GIP-1 levels were strongly reduced (Fig. 6C, D), similarly to NOCA-1. We also observed that PAR-6 depletion resulted in a decrease in the number of PTRN-1 puncta in the epidermis (Fig. 6E, F). As NOCA-1 is a direct interaction partner of PAR-6, we examined if the loss of GIP-1 is due to the loss of NOCA-1 localization. Indeed, in a *noca-1(ok3692)* deletion mutation GIP-1 levels were significantly reduced (Fig. 6G, H), suggesting that NOCA-1 acts upstream of GIP-1 in the *C. elegans* larval epidermis.

Finally, we examined if the failure to properly localize NOCA-1 could explain the microtubule defects we observed upon PAR-6 depletion. We determined microtubule bundle density, EB1 comet density, and microtubule growth rate in *noca-1(ok3692)* animals. In the *noca-1(ok3692)* deletion mutant, we observed a significant reduction in the density of circumferential microtubule bundles in the hypodermis (Fig. 5C, D). As reported in a previous study (Wang et al., 2015), we also observed reduced microtubule dynamics, with the density of growing MTs reduced by 65%, and microtubule growth rates reduced by 65 % (Fig. 5E, G). These values are all very similar to those we observed upon PAR-6 depletion and are consistent with a model in which the microtubule defects caused by PAR-6 depletion are a result of the requirement of PAR-6 in localizing NOCA-1. The effects on PTRN-1 may be a secondary consequence of microtubule defects caused by NOCA-1 loss.

## Discussion

Par6 and aPKC are essential for apical–basal polarization across animal species. Here, we used inducible protein degradation to identify an essential role for PAR-6 and PKC-3 in the larval epidermis of *C. elegans*, and a novel role for PAR-6 in regulating the assembly of microtubule bundles through its interaction partner NOCA-1/Ninein. Most studies of the apical PAR proteins in *C. elegans* have focused on embryonic tissues, and an essential role in postembryonic development has not been described. Depletion of *par-6, pkc-3*, or *par-3* by RNAi in larval stages caused defects in polarization of spermathecal cells and in ovulation, but not in larval development (Aono et al., 2004). Similar results were recently observed using a temperature sensitive *pkc-3* allele grown at non-permissive temperature (Montoyo-Rosario et al., 2020). More severe phenotypes were observed in hatching progeny (escapers) of *par-6, pkc-3*, or *par-3* RNAi-treated mothers, which showed partially penetrant defects in outgrowth of vulval precursor and seam cells, migrations of neuroblasts and axons, and the development of the somatic gonad (Welchman et al., 2007). The lack of a growth arrest phenotype in these studies presumably reflects incomplete gene inactivation.

### Auxin-inducible protein depletion of PAR proteins

The auxin-inducible degradation approach allowed us to bypass embryonic requirements and examine the roles of PAR-6, PKC-3, and PAR-3 in specific epithelial tissues during larval development. Despite these advantages, one drawback of all protein degradation approaches is that it remains difficult to draw conclusions from negative results. Although we tagged all known PAR-3 protein isoforms and observed efficient protein depletion, ubiquitous depletion of PAR-3 did not cause obvious defects in larvae. While PAR-3 may indeed not be essential in larval tissues, it is also possible that very low levels of PAR-3 are sufficient for its functioning, or that unpredicted splicing events cause the expression of non-degron tagged PAR-3 isoforms. One approach to counteract the latter possibility would be to replace the endogenous gene with a re-engineered copy that is unlikely to express alternative splice variants, e.g. by replacing natural introns with artificial ones and removing internal promoters. However, removing this level of regulation and expressing only one isoform may affect the functioning of *par-3* and cause unintended side effects. We also did not detect phenotypes upon depletion of PAR-6 or PKC-3 in the larval intestine. Similar caveats as for PAR-3 depletion apply here, though PAR-6 depletion did lead to complete loss of PKC-3 from the apical domain, and epidermal depletion caused severe phenotypes. These observations make it less likely that the lack of a phenotype is due to the expression of unknown isoforms. PAR-6 and PKC-3 are likely to play non-essential or redundant roles in the intestine, as a previous study found that PAR-6 contributes to endosome positioning in this tissue (Winter et al., 2012).

### Roles of PAR-6 and PKC-3 in junction formation and cell polarity

Depletion of PAR-6 and PKC-3 in the epidermis resulted in a fragmented appearance of the hyp7/seam and seam/seam junctions, similar to previous observations in embryonic epithelia (Montoyo-Rosario et al., 2020; Totong et al., 2007; Von Stetina and Mango, 2015; Von Stetina et al., 2017). Our data therefore further demonstrate a general requirement for PAR-6 and PKC-3 in junction formation or maintenance. We also examined if PKC-3 functions to exclude the basolateral polarity proteins PAR-1 and LGL-1 from the apical domain. In the epidermis, PKC-3 depletion caused a rapid invasion of LGL-1 in the apical domain of the seam cells, while PAR-1 remained junctional and basal. Thus, as in the one cell embryo, PKC-3 functions to exclude LGL-1 in the seam cells. A recent study found that LGL-1 can suppress sterility of a temperature sensitive *pkc-3* allele, further indicating that the interaction between LGL-1 and PKC-3 is functionally relevant (Montoyo-Rosario et al., 2020).

In contrast to the epidermis, LGL-1 localization in the intestine remained unchanged upon PKC-3 depletion, and we observed no obvious abnormalities in the intestine upon PAR-6 or PKC-3 depletion. Thus, while PAR-6 and PKC-3 are essential for development of the embryonic intestine (Totong et al., 2007), they do not appear to be essential in the larval intestine. Other cellular systems, such as polarized protein trafficking, may suffice to maintain cell polarity in the absence of the apical PAR proteins (Shafaq-Zadah et al., 2012; Zhang et al., 2011; Zhang et al., 2012). An analogous situation exists in the Drosophila midgut, where integrins, but not the apical PAR proteins, are essential for polarization (Chen et al., 2018). The lack of LGL-1 mislocalization also points to the existence of possible redundancies in polarization of cortical polarity regulators, which may be uncovered through enhancer screens in PAR-6 or PKC-3 depleted backgrounds.

In embryonic epithelia, the requirements of the apical PAR proteins also vary between tissues. Intestinal and epidermal cells depleted of PAR-6 or PKC-3 using the ZF1 system still show apicobasal polarization, as evidenced by apical localization of junctional and cytoskeletal proteins (Montoyo-Rosario et al., 2020; Totong et al., 2007). However, in the arcade cells of the pharynx, most PAR-6 depleted animals show no apical enrichment of junctional or apical cytoskeletal markers (Von Stetina and Mango, 2015). These data further highlight that the requirements for PAR-6 and PKC-3 can vary between tissues.

### Interdependencies between the PAR proteins

PAR-6 and PKC-3 were mutually dependent in both the epidermis and intestine. This result was not surprising, as Par6 and aPKC act as a dimer and have been shown to be mutually dependent in other *C. elegans* tissues (Hung and Kemphues, 1999; Li et al., 2010a; Nance et al., 2003; Tabuse et al., 1998; Totong et al., 2007). We also found that depletion of PAR-3 from the epidermis resulted in a partial loss of PAR-6. This is consistent with findings that multiple mechanisms can promote cortical recruitment of Par6–aPKC, including binding of Par6 to Par3 or Cdc42, and binding of aPKC to phospholipids (Dong et al., 2020; Hong et al., 2003; Hutterer et al., 2004; Joberty et al., 2000; Nagai-Tamai et al., 2002; Nunes de Almeida et al., 2019; Rodriguez et al., 2017; Wang et al., 2017; Wodarz et al., 2000). Conversely, the apical localization of PAR-3 was dependent upon PKC-3. While Par3 often localizes independently of Par6/aPKC, a similar dependency has been observed in Drosophila and in the *C. elegans* zygote (Hutterer et al., 2004; Petronczki and Knoblich, 2001; Tabuse et al., 1998; Watts et al., 1996).

Two results were more surprising in light of the interdependency between PAR-6 and PKC-3. First, whereas depletion of PKC-3 causes a loss of apical PAR-3, depletion of PAR-6 did not have a similar effect. Second, PAR-3 depletion resulted in a partial reduction of apical PAR-6 levels, but not of PKC-3. An independence in PAR-6 and PKC-3 localization has been observed previously in the one cell embryo, where reduction of the chaperone protein CDC-37 by RNAi resulted in the localization of PAR-6 to the cortex independently of PKC-3 (Beers and Kemphues, 2006). Thus, under certain conditions, PAR-6 and PKC-3 may not act as an obligate dimer.

### A novel role for PAR-6 in epidermal microtubule organization

Epidermal specific depletion uncovered a novel role for PAR-6 in organizing non-centrosomal microtubule bundles. In epithelial cells, apical non-centrosomal microtubule organizing centers (ncMTOCs) assemble apical–basal microtubule arrays. ncMTOCs contain proteins and complexes involved in microtubule anchoring, microtubule stabilization, and microtubule nucleation — such as the γ-tubulin ring complex (γ-TuRC) (Sanchez and Feldman, 2017). How apical ncMTOCs are organized is not well understood, but several studies indicate an important role for apical PAR proteins in this process. In the cellularizing Drosophila embryo, aPKC is required for the transition from centrosome emanated asters to non-centrosome associated apical–basal bundles (Harris and Peifer, 2007). In the developing embryonic intestine of *C. elegans*, PAR-3 is needed for the redistribution of γ-tubulin and other microtubule regulators from the centrosomes to the apical domain of the cell (Feldman and Priess, 2012). A role for Par6 in regulating microtubule organizing centers may not be limited to epithelial ncMTOCs. For example, in several mammalian cultured cell lines Par6 is a component of centrosomes and regulates centrosomal protein composition (Dormoy et al., 2013; Kodani et al., 2010).

Epidermal depletion of PAR-6 resulted in reduced numbers of circumferential microtubule bundles, as well as a reduced microtubule growth rate and EB1 comet density. Moreover, depletion of PAR-6 led to a loss of apical NOCA-1 enrichment at seam–seam and seam–hyp7 junctions. The effects of PAR-6 depletion on microtubule organization and dynamics are very similar to those we observed in a *noca-1* mutant. While other models are possible, these data are consistent with PAR-6 acting through NOCA-1 to control microtubule organization in the epidermis. The reduced microtubule growth rate and EB1 comet density we observed in *noca-1* mutant animals have been reported previously (Wang et al., 2015). However, no defects in circumferential microtubule bundle density were observed in that study, despite using the same *noca-1(ok3692)* allele. The observed difference may be a result of a difference in exact experimental procedure or the precise genetic background used. For example, whereas we used the microtubule-binding protein GFP::MAPH-1.1 to label microtubules, the study by Wang *et al.* used a GFP::β-tubulin fusion.

We also found that, in the epidermis, the localization of GIP-1 is dependent on NOCA-1. The relationship between NOCA-1 and γ-TuRC components has been examined previously in two different tissues (Sallee et al., 2018; Wang et al., 2015). In the germline, NOCA-1 co-localizes with γ-tubulin to non-centrosomal microtubule arrays but is not required for the localization of γ-tubulin (Wang et al., 2015). In fact, in this tissue the localization of a short NOCA-1 protein lacking isoform-specific N-terminal extensions is dependent for its localization on γ-tubulin. The longer NOCA-1h isoform, however, localizes independently of γ-tubulin, indicating the presence of multiple NOCA-1 localization signals (Wang et al., 2015). In the embryonic intestine, the localization of NOCA-1 was not altered by the depletion of GIP-1 (Sallee et al., 2018). However, microtubule organization in the intestine is regulated differently from the epidermis, as apical microtubule organization was largely normal even in *ptrn-1* mutant animals depleted of intestinal NOCA-1 and GIP-1 (Sallee et al., 2018). Thus, differential effects of γ-TuRC component loss may reflect differences in the mechanisms of microtubule regulation. Whether PAR-6 plays a role in ncMTOC assembly and microtubule organization in tissues other than the epidermis remains to be investigated.

In addition to the effects on NOCA-1 and GIP-1, PAR-6 depletion resulted in a reduced number of PTRN-1 puncta in the epidermis. PTRN-1 is a member of the Patronin/CAMSAP/Nezha family of minus end-associated proteins, which stabilize and protect uncapped microtubule minus ends (Atherton et al., 2019; Goodwin and Vale, 2010; Hendershott and Vale, 2014; Jiang et al., 2014). NOCA-1 was previously shown to act in parallel with PTRN-1 in organizing circumferential microtubule arrays in the *C. elegans* epidermis (Wang et al., 2015). The mechanistic details of the relationship between NOCA-1 and PTRN-1 have not been resolved, but their distinct localization patterns suggest that they act on distinct pools of microtubules. Our data does not reveal why PAR-6 depletion results in a reduced number of PTRN-1 foci, but a likely possibility is that this is a secondary consequence of the microtubule defects caused by the loss of NOCA-1 localization.

### Mechanisms of larval growth arrest

The depletion of PAR-6 or PKC-3 in the epidermis led to a rapid growth arrest and cessation of the molting cycle. What causes this dramatic effect on animal development? The junctional defects we observed are unlikely to be the primary consequence, as effects on cell junctions appeared only after 24 h of exposure to auxin. The effects on LGL-1 were more rapid but are also not likely to explain the growth arrest, as *lgl-1* mutants are viable (Beatty et al., 2010; Hoege et al., 2010). The effects on the microtubule cytoskeleton are likely to contribute to the growth arrest or molting defects. However, *noca-1* mutants displayed similar microtubule defects as PAR-6 depletion yet develop to adulthood. Interestingly, *noca-1; ptrn-1* double mutant animals do grow slowly and frequently die before reaching adulthood (Wang et al., 2015). Thus, the combined defects in NOCA-1 and PTRN-1 localization we observed upon PAR-6 depletion may partially explain the growth defects. The roles of PTRN-1 may not be limited to microtubule regulation, as a recent study demonstrated that PTRN-1 stimulates actin polymerization during endocytic recycling in the intestine (Gong et al., 2018). A final possibility is that the molting defect is the primary cause of the developmental arrest, as failure to molt can cause a growth arrest (Brooks et al., 2003; Lažetić and Fay, 2017; Russel et al., 2011; Yochem et al., 1999).

One way in which PAR-6 and PKC-3 could affect molting is by affecting intracellular trafficking. Molting requires the coordinated activity of the endocytic and exocytic machineries to mediate apical secretion of new cuticle components, recycling of old cuticle components, and uptake of sterols from the environment (Lažetić and Fay, 2017). Accordingly, mutations in multiple regulators of intracellular trafficking cause molting defects (Lažetić and Fay, 2017). Several links between cortical polarity regulators and the polarized trafficking machinery have been uncovered (Rodriguez-Boulan and Macara, 2014). In *C. elegans*, *par-3, par-6*, and *pkc-3* were all found to be required for endocytic trafficking in oocytes, and RNAi for *par-3* and *par-6* causes scattering of multiple endosome types in the intestine (Balklava et al., 2007; Winter et al., 2012). It is possible, therefore, that PAR-6 and PKC-3 regulate vesicle trafficking in molting as well. Such regulation may be indirect, through regulation of cytoskeletal components, or through more direct mechanisms remaining to be uncovered.

In summary, our data supports that PAR-6 and PKC-3 have multiple roles in the epidermis that support larval development and molting. We have also uncovered an important role for PAR-6 in regulating the microtubule cytoskeleton, while additional mechanisms through which PAR-6 and PKC-3 control growth and/or molting likely remain to be discovered.

## Materials and Methods

### *C. elegans* strains

All *C. elegans* strains used in this study are derived from the N2 Bristol strain, and are listed in Table 1. All strains were maintained at 20 °C on Nematode Growth Medium (NGM) plates seeded with *Escherichiae coli* OP50 bacteria under standard conditions (Brenner, 1974).

**Table 1.**
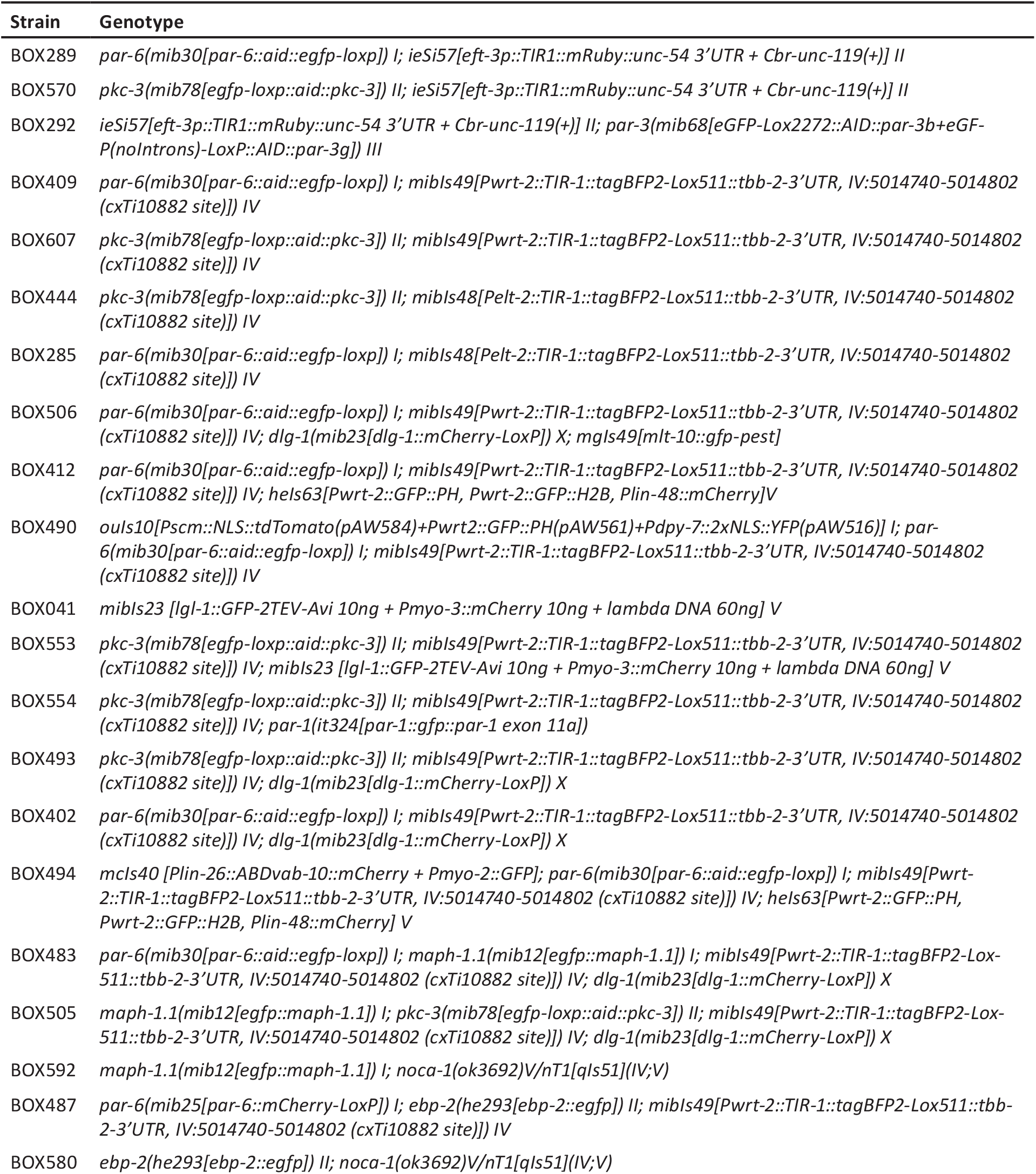
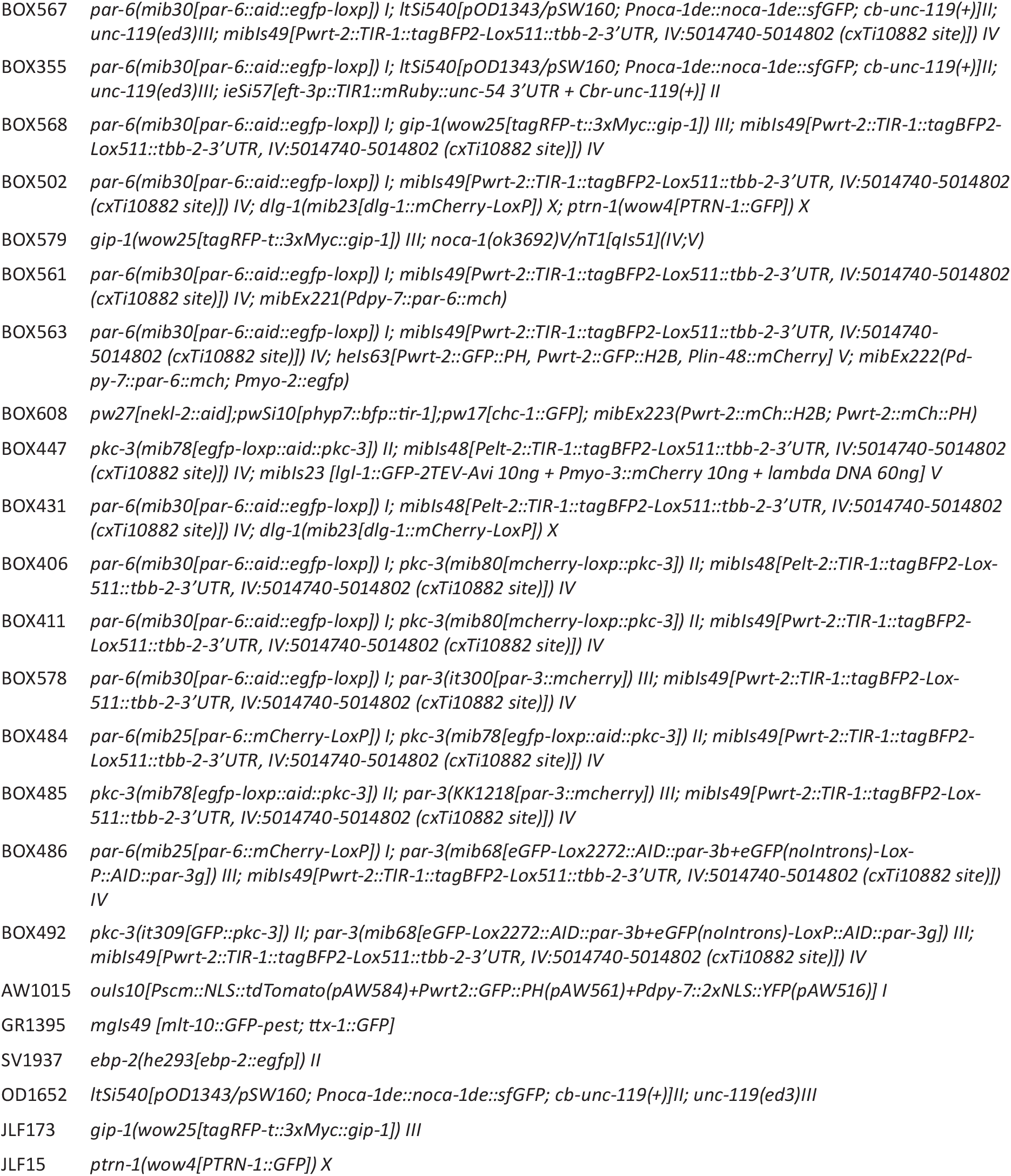
List of strains used

### CRISPR/Cas9 genome engineering

All gene editing was done by homology-directed repair of CRISPR/Cas9-induced DNA double-strand breaks, using plasmid-based expression of Cas9 and sgRNAs. All edits were made in an N2 background, with the exception of 2x*(egfp::aid)::par-3*, for which *egfp::aid::par-3* was used as the starting background. All fusions were repaired using a plasmid-based template with 190–600 bp homology arms and containing a self-excising cassette (SEC) for selection (Dickinson et al., 2015). The homology arms included mutations of the sgRNA recognition sites to prevent re-cutting after repair. The *par-6::aid::egfp, par-6::mCherry, dlg-1::mCherry* and *ebp-2::egfp* vectors were cloned using Gibson assembly and vector pJJR82 (Addgene #75027) (Gibson et al., 2009; Ramalho et al., 2020) as the backbone. The *2x(egfp::aid)::par-3*, *Pwrt-2::tir-1::bfp* and *Pelt-2::tir-1::bfp* vectors were cloned using SapTrap assembly into vector pMLS257 (Schwartz and Jorgensen, 2016), and the *egfp::aid::pkc-3* and *mCherry::pkc-3* vectors were cloned using SapTrap assembly into vector pDD379 (Dickinson et al., 2018). The sgRNAs were expressed from a plasmid under control of a U6 promoter. To generate sgRNA vectors, antisense oligonucleotide pairs were annealed and ligated into BbsI-linearized pJJR50 (Addgene #75026) (Waaijers et al., 2016), with the exception of the *pkc-3* fusions, in which the sgRNA was incorporated into assembly vector pDD379 using SapTrap assembly. The targeted sequences can be found in Table 2. Injection mixes were prepared in MilliQ H_2_ O and contained 50 ng/ml *Peft-3::cas9* (Addgene ID #46168) (Friedland et al., 2013), 50–100 ng/μl *U6::sgRNA*, 50 ng/μl of repair template, with the exception of the *pkc-3* fusions, in which the sgRNA-repair-template vector was used at a concentration of 65 ng/μl. All mixes also contained 2.5 ng/μl of the co-injection pharyngeal marker *Pmyo-2::GFP* or *Pmyo-2::tdTomato* to aid in visual selection of transgenic strains. Young adult hermaphrodites were injected in the germline using an inverted micro-injection setup (Eppendorf FemtoJet 4x mounted on a Zeiss Axio Observer A.1 equipped with an Eppendorf Transferman 4r). Candidate edited progeny were selected on plates containing 250 ng/μl of hygromycin (Dickinson et al., 2015), and correct genome editing was confirmed by Sanger sequencing (Macrogen Europe) of PCR amplicons encompassing the edited genomic region. From correctly edited strains, the hygromycin selection cassette was excised by a heat shock of L1 larvae at 34 °C for 1 h in a water bath. Correct excision was confirmed by Sanger sequencing. Sequence files of the final gene fusions in Genbank format are in Supplementary File 1.

**Table 2.**
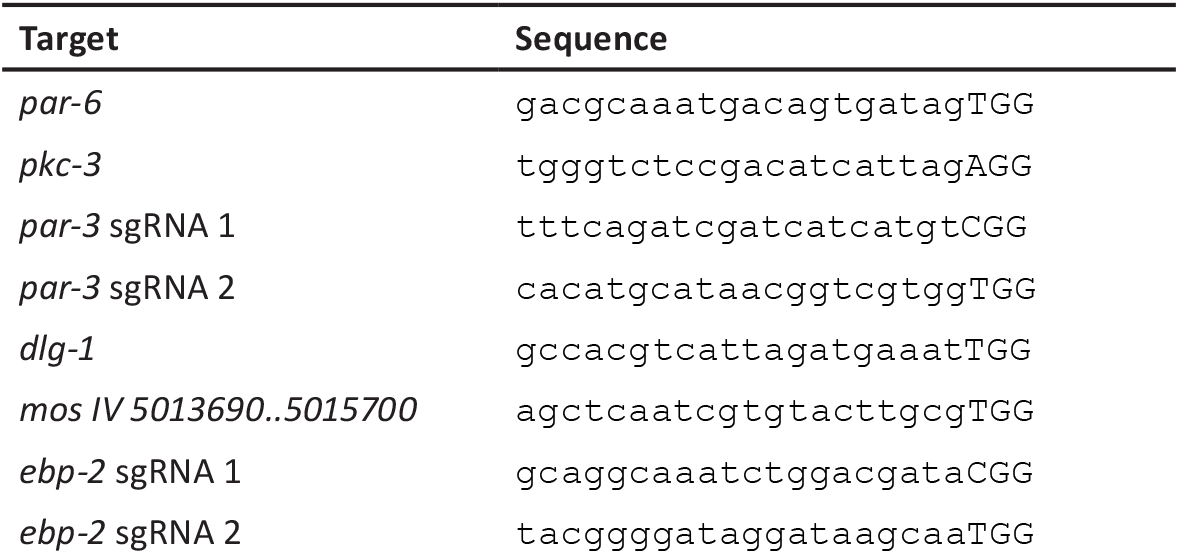
List of sgRNA target sites

### *C. elegans* synchronization

In order to obtain synchronized worm populations, plates with eggs were carefully washed with M9 (0.22 M KH_2_PO_4_, 0.42 M Na_2_HPO_4_, 0.85 M NaCl, 0.001 M MgSO_4_) buffer in order to remove larvae and adults but leave the eggs behind. Plates were washed again using M9 buffer after an hour, to collect larvae hatched within that time span.

### Auxin Inducible Degradation

Auxin treatment was performed by placing synchronized populations of worms on NGP plates seeded with *E. coli* OP50 and containing 1 or 4 mM auxin. To prepare plates, auxin (Alfa Aesar A10556) was diluted into the autoclaved NGM agar medium after cooling to 60 °C prior to plate pouring. Plates were kept for a maximum of 2 weeks in the dark at 4 °C.

### *C. elegans* growth curves

To measure growth curves, L1 animals synchronized as described above were placed on NGM plates seeded with *E. coli* OP50 and either lacking auxin or containing 4 mM auxin. Images were taken in 24 h intervals up to 96 h, using a Zeiss Axio Zoom.V16 equipped with a PlanNeoFluar Z 1x/0.25 objective and Axiocam 506 color camera, driven by Zen Pro software. Animal length was quantified in ImageJ(FIJI) software by drawing a spline along the center line of the animal (Rueden et al., 2017; Schindelin et al., 2012).

### Feeding RNAi

*pak-1* RNAi clones were obtained from the genome-wide Ahringer and Vidal RNAi feeding librarys (Kamath and Ahringer, 2003; Rual et al., 2004). Ahringer clone supplied through Source BioScience. An HT115 clone expressing the L4440 vector lacking an insert was used as a control. For feeding RNAi experiments, bacteria were cultured overnight in 10 mL of Lysogeny Broth (LB) supplemented with 100 μg/ml ampicilin (Amp) and 2.5 μg/ml tetracyclin (Tet) at 37°C in an incubator rotating at 200 rpm. Cultures were incubated for 120 min in the presence of 1 mM Isopropyl β-D-1-thiogalactopyranoside (IPTG) to induce production of dsRNA. Cultures were pelleted by centrifugation at 4000 g for 15 min and concentrated 5x. NGM agar plates supplemented with 50 μg/ml Amp, 12.5 μg/mL Tet and 1 mM IPTG were seeded with 250 μl of bacterial suspension. 8-10 L4 hermaphrodites per strain were transferred to NGM-RNAi plates, and two plates were analyzed per condition. The F1 generation was synchronized and its growth curved determined as described above.

### Molting assay

Synchronized L1 animals were placed on NGM plates seeded with *E. coli* OP50 and either lacking auxin or containing 1 mM auxin. Fluorescence images were taken in 1 h intervals from 11 h to 32 h of development, using a Zeiss Axio Zoom. V16 equipped with a PlanNeoFluar Z 1x/0.25 objective and Axiocam 506 color camera, driven by Zen Pro software. Expression levels of the *Pmlt-10::gfp::pest* reporter were quantified in ImageJ(FIJI) software (see image analysis).

### Seam cell lineage analysis

Synchronized animals were placed on NGM plates seeded with *E. coli* OP50 and either lacking auxin or containing 1 mM auxin. Animals were placed on plates immediately after hatching (L1 degradation), at 7 h of development (L2 degradation) or at 19 h of development (L3 degradation). Every hour, from placing the worms on plates until seam cell fusion on control plates lacking auxin, a sample of animals were imaged by spinning disc confocal microscopy. The number of seam cell and hyp7 nuclei were counted manually.

### Microscopy

Live imaging of *C. elegans* larvae was done by mounting larvae on 5% agarose pads in a 10 mM Tetramisole solution in M9 buffer to induce paralysis. DIC imaging was performed with an upright Zeiss AxioImager Z2 microscope using a 63×1.4 NA objective and a Zeiss AxioCam 503 monochrome camera, driven by Zeiss Zen software. Spinning disk confocal imaging was performed using a Nikon Ti-U microscope driven by MetaMorph Microscopy Automation & Image Analysis Software (Molecular Devices) and equipped with a Yokogawa CSU-X1-M1 confocal head and an Andor iXon DU-885 camera, using 60× or 100× 1.4 NA objectives. All stacks along the z-axis were obtained at 0.25 μm intervals, and all images were analyzed and processed using ImageJ(FIJI) and Adobe Photoshop. For quantifications, the same laser power and exposure times were used within experiments.

### Quantitative image analysis

All image analysis was done in using ImageJ (FIJI). For intensity profile measurements of spinning disk microscopy data, background values were subtracted from the intensity measurements. Mean background intensity was quantified on a circular region in an area not containing any animals, except in quantifications in Fig. 4A, 4C and 6A, where background intensity was quantified on a circular region in an area with no fluorescence inside the worm.

For the intensity profiles in the epidermis, except those of RFP::GIP-1, a 10 px-wide line was drawn in the apical focal plane, from the hyp7 cytoplasm to the seam cell cytoplasm. For the intensity profiles of RFP::GIP-1, 10 20-px wide lines were drawn per cell in the apical focal plane, from the hyp7 cytoplasm to the seam cell cytoplasm, and then averaged to obtain a single intensity profile per cell. For the intensity profiles in the intestine, 8 50 px-wide lines were drawn from the lumen to the cytoplasm of the intestinal cells and averaged to obtain a single value per worm. The intensity profiles were manually aligned at the apical peak value.

To quantify the fluorescence intensity for the molting assay, whole worm fluorescence was quantified. An ROI of each whole worm was created by drawing a freehand line around the worm using the transmitted light channel. The corresponding fluorescence of the ROI was measured in the GFP channel.

Microtubule bundles were counted manually. A 5-px-wide freehand line was drawn along the worm at the dorsal/ ventral region, and the intensity profile was obtained. The number of fluorescent peaks was counted, and the microtubule bundle density was calculated by dividing the number of peaks by the measured distance.

EBP-2::GFP comet counting was done manually. Comet density was calculated by dividing the number of EBP-2::GFP comets by the surface of the area analyzed. Comets in both the lateral and dorsal/ventral hypodermis were measured.

PTRN-1::GFP puncta counting was done manually. Puncta density was calculated by dividing the number of PTRN-1::GFP puncta by the surface of the area analyzed. Puncta in both the seam cells and the hypodermis were measured.

Microtubule growth rate was calculated in an automated manner using the ImageJ plug-in ‘TrackMate’ (Tinevez et al., 2017). The following parameters were chosen: estimated blob diameter = 0.700 um; threshold = 200,000; simple LAP tracker; linking max distance = 1.5 um; gap-closing max distance = 1.5 um; gap-closing max frame gap = 3; duration of track = 10. The mean speed of the comets was averaged to obtain the average microtubule growth rate. Comets in both the seam cells and the hypodermis were measured.

To determine the directionality of the actin bundles and microtubule growth, images or movies were rotated to orient the seam cells horizontally. Lines were drawn along the microtubule or actin bundles, and the angle of these lines was calculated relative to the horizontal axis. Movies of EBP-2 were used to calculate the directionality of microtubule growth, where the direction of growth of individual comets was annotated manually. Maximum projections of EBP-2 movies were used to calculate the directionality of microtubule growth. Rose plots were generated using MatLAB.

### Seam lineage analysis

To generate the seam cell lineage the number of seam cells and hyp7 nuclei were analyzed at 1 h intervals, using the marker *ouIs10[scmp::NLS::tdTomato; dpy-7p::2xNLS::YFP;wrt-2p::GFP::PH]* that marks the seam nuclei in red and the hypodermal nuclei in green. Animals were classified according to showing a wild-type seam cell division pattern, having developmental defects such as delayed or arrested seam cell divisions, or having inappropriate seam cell differentiation. Control animals were classified at each larval stage. PAR-6 depleted animals were classified after they had undergone the delayed L2-stage divisions. From the total number of worms analyzed, the percentages of worms in each category were calculated.

### PAR-6::mCherry transgenic array

The *Pdpy-7::par-6::mCherry* vector used for PAR-6 hypodermal rescue was cloned into the pBSK(+) vector using Gibson assembly. The promoter of *dpy7*, which is expressed in hyp7 but not in the seam cells (Gilleard et al., 1997; Myers and Greenwald, 2005), was amplified from *C. elegans* genomic DNA using primers 5’-TGTAATACGACTCACTATAGGGCGAATTGGctcattccacgatttctcgc and 5’-tctggaacaaaatgtaagaatattc. A fragment of 5.3 kb containing the entire genomic sequence of *par-6* and a fragment of 402 bp of the *par-6* 3’ UTR were amplified from *C. elegans* N2 genomic DNA using primers 5’-tttaagaatattcttacattttgttccagaATGTCCTACAACGGCTCCTA and 5’-GGCCATGTTGTCCTCCTCTCCCTTGGACATGTCCTCTCCACTGTCCGAAT, and 5’-CACTCCACCGGAGGAATGGACGAGCTCTACTGAaaaactcttttcagcca and 5’-TAAAGGGAACAAAAGCTGGAGCTCCACCGCgaaataaataatttattctc, respectively. mCherry was amplified from pJJR83 (Addgene #75028) using primers 5’-TCCAAGGGAGAGGAGGACAA and 5’-GTAGAGCTCGTCCATTCCTC. Correct amplification and assembly was confirmed by Sanger sequencing. The plasmid generated can be found in Supplementary File 1. To generate transgenic lines young adult hermaphrodites were injected in the germline with 30 ng/μl of *Pdpy-7::par-6::mCherry*. mCherry fluorescence was used to select stable transgenic lines.

### Yeast two-hybrid analysis

Sequences encoding the PAR-6 PDZ domain and full-length NOCA-1d were PCR amplified from a mixed-stage cDNA library using primers par-6_F: 5’-ggaggcgcgccATGATTGTGCCAGAAGCTCATCG, par-6_R: 5’-ggagcggccgcTCAGGCGTTCGGTGTTCCTTGTT, noca-1d_F: 5’-ggaggcgcgccATGAATATTTGTTGTTGTGG and noca-1d_R: 5’-ggagcggccgcCTATTGAACTCTGCATACAT. PCR products were digested with AscI and NotI, and cloned into Gal4-DB vector pMB28 and Gal4-AD vector pMB29, respectively (Koorman et al., 2016). The resulting plasmids were transformed into *Saccharomyces cerevisiae* strains Y8930 (MATα) and Y8800 (MAT**a**) (Yu et al., 2008) using the Te/ LiAc transformation method (Schiestl and Gietz, 1989). DB::PAR-6/AD::NOCA-1 diploid yeast was generated by mating, and plated on synthetic defined (SD) medium plates lacking leucine, tryptophan, and histidine containing 2 mM 3-Amino-1,2,4-triazole (3-AT); and lacking leucine, tryptophan, and adenine to assess the presence of an interaction, and on an SD plate lacking leucine and histidine containing 1 mg/ml cycloheximide to test for self-activation by the DB::PAR-6 plasmid in the absence of the AD::NOCA-1 plasmid. Controls of known reporter activition strength and behavior on cycloheximide were also added to all plates.

### Statistical analysis

All statistical analyses were performed using GraphPad Prism 8. For population comparisons, a D’Agostino & Pearson test of normality was first performed to determine if the data was sampled from a Gaussian distribution. For data drawn from a Gaussian distribution, comparisons between two populations were done using an unpaired t test, with Welch’s correction if the SDs of the populations differ significantly, and comparisons between >2 populations were done using a one-way ANOVA, or a Welch’s ANOVA if the SDs of the populations differ significantly. For data not drawn from a Gaussian distribution, a non-parametric test was used (Mann-Whitney for 2 populations and Kruskal-Wallis for >2 populations). ANOVA and non-parametric tests were followed up with multiple comparison tests of significance (Dunnett’s, Tukey’s, Dunnett’s T3 or Dunn’s). Tests of significance used and sample sizes are indicated in the figure legends. No statistical method was used to pre-determine sample sizes. No samples or animals were excluded from analysis. The experiments were not randomized, and the investigators were not blinded to allocation during experiments and outcome assessment.

## Supporting information

Supplemental DNA files

## Acknowledgements

We thank R. Schmidt, A. Homavar and S. van den Heuvel for sharing strain SV1937, J. Feldman for strains JLF15 and JLF173, K. Oegema for strain OD1652, A. Woollard for strain AW1015, and A. Frand for strain GR1395. We also thank D. Fay for distributing some of these strains. We thank S. van den Heuvel, M. Harterink, D. Fay and members of the S. van den Heuvel and M. Boxem groups for helpful discussions, M. Harterink for critical reading of the manuscript, and J. Cravo for generating the rose plots. We also thank Wormbase (Harris et al., 2020) and the Biology Imaging Center, Faculty of Sciences, Department of Biology, Utrecht University. Some strains were provided by the Caenorhabditis Genetics Center, which is funded by NIH Office of Research Infrastructure Programs (P40 OD010440).

## Author contributions

Conceptualization: V.G.G., H.R.P., M.B.; Methodology: V.G.G., H.R.P.; Formal analysis: V.G.G., H.R.P.; Investigation: R.R.B., A.R., J.K.; Writing-original draft: V.G.G., H.R.P., M.B.; Visualization: V.G.G., H.R.P.; Supervision: M.B.; Project administration: M.B.; Funding acquisition: M.B.

## Supplementary materials

This manuscript is accompanied by three supplementary material files: Supplementary Figures.pdf, Supplemental File 1 - DNA files.zip, and Video S1.m4v.

## References

Aceto, D., Beers, M. and Kemphues, K. J.(2006). Interaction of PAR-6 with CDC-42 is required for maintenance but not establishment of PAR asymmetry in C. elegans. Dev. Biol. 299, 386–397.

Achilleos, A., Wehman, A. M. and Nance, J.(2010). PAR-3 mediates the initial clustering and apical localization of junction and polarity proteins during C. elegans intestinal epithelial cell polarization. Dev. Camb. Engl. 137, 1833–1842.

Aguilar-Aragon, M., Elbediwy, A., Foglizzo, V., Fletcher, G. C., Li, V. S. W. and Thompson, B. J.(2018). Pak1 Kinase Maintains Apical Membrane Identity in Epithelia. Cell Rep. 22, 1639–1646.

Aono, S., Legouis, R., Hoose, W. A. and Kemphues, K. J.(2004). PAR-3 is required for epithelial cell polarity in the distal spermatheca of C. elegans. Dev. Camb. Engl. 131, 2865–2874.

Atherton, J., Luo, Y., Xiang, S., Yang, C., Rai, A., Jiang, K., Stangier, M., Vemu, A., Cook, A. D., Wang, S., et al.(2019). Structural determinants of microtubule minus end preference in CAMSAP CKK domains. Nat. Commun. 10, 5236.

Balklava, Z., Pant, S., Fares, H. and Grant, B. D.(2007). Genome-wide analysis identifies a general requirement for polarity proteins in endocytic traffic. Nat. Cell Biol. 9, 1066–1073.

Beatty, A., Morton, D. and Kemphues, K.(2010). The C. elegans homolog of Drosophila Lethal giant larvae functions redundantly with PAR-2 to maintain polarity in the early embryo. Development 137, 3995–4004.

Beers, M. and Kemphues, K.(2006). Depletion of the co-chaperone CDC-37 reveals two modes of PAR-6 cortical association in C. elegans embryos. Dev. Camb. Engl. 133, 3745–3754.

Betschinger, J., Mechtler, K. and Knoblich, J. A.(2003). The Par complex directs asymmetric cell division by phosphorylating the cytoskeletal protein Lgl. Nature 422, 326–330.

Bilder, D., Schober, M. and Perrimon, N.(2003). Integrated activity of PDZ protein complexes regulates epithelial polarity. Nat. Cell Biol. 5, 53–58.

Bossinger, O., Klebes, A., Segbert, C., Theres, C. and Knust, E.(2001). Zonula adherens formation in Caenorhabditis elegans requires dlg-1, the homologue of the Drosophila gene discs large. Dev Biol 230, 29–42.

Boxem, M., Maliga, Z., Klitgord, N., Li, N., Lemmens, I., Mana, M., de Lichtervelde, L., Mul, J. D., van de Peut, D., Devos, M., et al.(2008). A protein domain-based interactome network for *C. elegans* early embryogenesis. Cell 134, 534–545.

Brajenovic, M., Joberty, G., Küster, B., Bouwmeester, T. and Drewes, G.(2004). Comprehensive proteomic analysis of human Par protein complexes reveals an interconnected protein network. J. Biol. Chem. 279, 12804–12811.

Brenner, S.(1974). The Genetics of Caenorhabditis Elegans. Genetics 77, 71–94.

Brooks, D. R., Appleford, P. J., Murray, L. and Isaac, R. E.(2003). An essential role in molting and morphogenesis of Caenorhabditis elegans for ACN-1, a novel member of the angiotensin-converting enzyme family that lacks a metallopeptidase active site. J. Biol. Chem. 278, 52340–52346.

Chen, X. and Macara, I. G.(2005). Par-3 controls tight junction assembly through the Rac exchange factor Tiam1. Nat. Cell Biol. 7, 262–269.

Chen, J., Sayadian, A.-C., Lowe, N., Lovegrove, H. E. and St Johnston, D.(2018). An alternative mode of epithelial polarity in the Drosophila midgut. PLOS Biol. 16, e3000041.

Chisholm, A. D. and Hsiao, T. I.(2012). The Caenorhabditis elegans epidermis as a model skin. I: development, patterning, and growth. Wiley Interdiscip. Rev. Dev. Biol. 1, 861–878.

Chisholm, A. D. and Xu, S.(2012). The Caenorhabditis elegans epidermis as a model skin. II: differentiation and physiological roles. Wiley Interdiscip. Rev. Dev. Biol. 1, 879–902.

Chuang, M., Hsiao, T. I., Tong, A., Xu, S. and Chisholm, A. D.(2016). DAPK interacts with Patronin and the microtubule cytoskeleton in epidermal development and wound repair. eLife 5,.

Costa, M., Draper, B. W. and Priess, J. R.(1997). The role of actin filaments in patterning the Caenorhabditis elegans cuticle. Dev. Biol. 184, 373–384.

Dickinson, D. J., Pani, A. M., Heppert, J. K., Higgins, C. D. and Goldstein, B.(2015). Streamlined Genome Engineering with a Self-Excising Drug Selection Cassette. Genetics 200, 1035–1049.

Dickinson, D. J., Slabodnick, M. M., Chen, A. H. and Goldstein, B.(2018). SapTrap assembly of repair templates for Cas9-triggered homologous recombination with a self-excising cassette. MicroPublication Biol. 2018,.

Doerflinger, H., Vogt, N., Torres, I. L., Mirouse, V., Koch, I., Nüsslein-Volhard, C. and St Johnston, D.(2010). Bazooka is required for polarisation of the Drosophila anterior-posterior axis. Dev. Camb. Engl. 137, 1765–1773.

Dong, W., Lu, J., Zhang, X., Wu, Y., Lettieri, K., Hammond, G. R. and Hong, Y.(2020). A polybasic domain in aPKC mediates Par6-dependent control of membrane targeting and kinase activity. J. Cell Biol. 219,.

Dormoy, V., Tormanen, K. and Sütterlin, C.(2013). Par6γ is at the mother centriole and controls centrosomal protein composition through a Par6α-dependent pathway. J. Cell Sci. 126, 860–870.

Feldman, J. L. and Priess, J. R.(2012). A role for the centrosome and PAR-3 in the hand-off of MTOC function during epithelial polarization. Curr. Biol. CB 22, 575–582.

Feng, W., Wu, H., Chan, L.-N. and Zhang, M.(2008). Par-3-mediated junctional localization of the lipid phosphatase PTEN is required for cell polarity establishment. J. Biol. Chem. 283, 23440–23449.

Franz, A. and Riechmann, V.(2010). Stepwise polarisation of the Drosophila follicular epithelium. Dev. Biol. 338, 136–147.

Gally, C., Wissler, F., Zahreddine, H., Quintin, S., Landmann, F. and Labouesse, M.(2009). Myosin II regulation during C. elegans embryonic elongation: LET-502/ROCK, MRCK-1 and PAK-1, three kinases with different roles. Dev. Camb. Engl. 136, 3109–3119.

Georgiou, M., Marinari, E., Burden, J. and Baum, B.(2008). Cdc42, Par6, and aPKC regulate Arp2/3-mediated endocytosis to control local adherens junction stability. Curr. Biol. CB 18, 1631–1638.

Gibson, D. G., Young, L., Chuang, R.-Y., Venter, J. C., Hutchison, C. A. and Smith, H. O.(2009). Enzymatic assembly of DNA molecules up to several hundred kilobases. Nat. Methods 6, 343–345.

Gilleard, J. S., Barry, J. D. and Johnstone, I. L.(1997). cis regulatory requirements for hypodermal cell-specific expression of the Caenorhabditis elegans cuticle collagen gene dpy-7. Mol. Cell. Biol. 17, 2301–2311.

Giot, L., Bader, J. S., Brouwer, C., Chaudhuri, A., Kuang, B., Li, Y., Hao, Y. L., Ooi, C. E., Godwin, B., Vitols, E., et al.(2003). A protein interaction map of *Drosophila melanogaster*. Science 302, 1727–1736.

Goldstein, B. and Macara, I. G.(2007). The PAR proteins: fundamental players in animal cell polarization. Dev. Cell 13, 609–622.

Gong, T., Yan, Y., Zhang, J., Liu, S., Liu, H., Gao, J., Zhou, X., Chen, J. and Shi, A.(2018). PTRN-1/CAMSAP promotes CYK-1/formin-dependent actin polymerization during endocytic recycling. EMBO J. 37,.

Goodwin, S. S. and Vale, R. D.(2010). Patronin regulates the microtubule network by protecting microtubule minus ends. Cell 143, 263–274.

Green, R. A., Kao, H.-L., Audhya, A., Arur, S., Mayers, J. R., Fridolfsson, H. N., Schulman, M., Schloissnig, S., Niessen, S., Laband, K., et al.(2011). A High-Resolution C. elegans Essential Gene Network Based on Phenotypic Profiling of a Complex Tissue. Cell 145, 470–482.

Grossmann, A., Benlasfer, N., Birth, P., Hegele, A., Wachsmuth, F., Apelt, L. and Stelzl, U.(2015). Phospho-tyrosine dependent protein-protein interaction network. Mol. Syst. Biol. 11, 794.

Harris, T. J. C. and Peifer, M.(2004). Adherens junction-dependent and -independent steps in the establishment of epithelial cell polarity in Drosophila. J. Cell Biol. 167, 135–147.

Harris, T. J. C. and Peifer, M.(2005). The positioning and segregation of apical cues during epithelial polarity establishment in Drosophila. J. Cell Biol. 170, 813–823.

Harris, T. J. C. and Peifer, M.(2007). aPKC controls microtubule organization to balance adherens junction symmetry and planar polarity during development. Dev. Cell 12, 727–738.

Harris, K. P. and Tepass, U.(2008). Cdc42 and Par proteins stabilize dynamic adherens junctions in the Drosophila neuroectoderm through regulation of apical endocytosis. J. Cell Biol. 183, 1129–1143.

Harris, T. W., Arnaboldi, V., Cain, S., Chan, J., Chen, W. J., Cho, J., Davis, P., Gao, S., Grove, C. A., Kishore, R., et al.(2020). WormBase: a modern Model Organism Information Resource. Nucleic Acids Res. 48, D762–D767.

Hein, M. Y., Hubner, N. C., Poser, I., Cox, J., Nagaraj, N., Toyoda, Y., Gak, I. A., Weisswange, I., Mansfeld, J., Buchholz, F., et al.(2015). A human interactome in three quantitative dimensions organized by stoichiometries and abundances. Cell 163, 712–723.

Hendershott, M. C. and Vale, R. D.(2014). Regulation of microtubule minus-end dynamics by CAMSAPs and Patronin. Proc. Natl. Acad. Sci. U. S. A. 111, 5860–5865.

Hirano, Y., Yoshinaga, S., Takeya, R., Suzuki, N. N., Horiuchi, M., Kohjima, M., Sumimoto, H. and Inagaki, F.(2005). Structure of a cell polarity regulator, a complex between atypical PKC and Par6 PB1 domains. J. Biol. Chem. 280, 9653–9661.

Hoege, C., Constantinescu, A.-T., Schwager, A., Goehring, N. W., Kumar, P. and Hyman, A. A.(2010). LGL can partition the cortex of one-cell Caenorhabditis elegans embryos into two domains. Curr. Biol. 20, 1296–1303.

Hong, Y., Ackerman, L., Jan, L. Y. and Jan, Y.-N.(2003). Distinct roles of Bazooka and Stardust in the specification of Drosophila photoreceptor membrane architecture. Proc. Natl. Acad. Sci. U. S. A. 100, 12712–12717.

Hung, T. J. and Kemphues, K. J.(1999). PAR-6 is a conserved PDZ domain-containing protein that colocalizes with PAR-3 in Caenorhabditis elegans embryos. Dev. Camb. Engl. 126, 127–135.

Hurov, J. B., Watkins, J. L. and Piwnica-Worms, H.(2004). Atypical PKC phosphorylates PAR-1 kinases to regulate localization and activity. Curr. Biol. CB 14, 736–741.

Hutterer, A., Betschinger, J., Petronczki, M. and Knoblich, J. A.(2004). Sequential roles of Cdc42, Par-6, aPKC, and Lgl in the establishment of epithelial polarity during Drosophila embryogenesis. Dev. Cell 6, 845–854.

Huttlin, E. L., Ting, L., Bruckner, R. J., Gebreab, F., Gygi, M. P., Szpyt, J., Tam, S., Zarraga, G., Colby, G., Baltier, K., et al.(2015). The BioPlex Network: A Systematic Exploration of the Human Interactome. Cell 162, 425–440.

Izumi, Y., Hirose, T., Tamai, Y., Hirai, S., Nagashima, Y., Fujimoto, T., Tabuse, Y., Kemphues, K. J. and Ohno, S.(1998). An atypical PKC directly associates and colocalizes at the epithelial tight junction with ASIP, a mammalian homologue of Caenorhabditis elegans polarity protein PAR-3. J. Cell Biol. 143, 95–106.

Jiang, K., Hua, S., Mohan, R., Grigoriev, I., Yau, K. W., Liu, Q., Katrukha, E. A., Altelaar, A. F. M., Heck, A. J. R., Hoogenraad, C. C., et al.(2014). Microtubule minus-end stabilization by polymerization-driven CAMSAP deposition. Dev. Cell 28, 295–309.

Joberty, G., Petersen, C., Gao, L. and Macara, I. G.(2000). The cell-polarity protein Par6 links Par3 and atypical protein kinase C to Cdc42. Nat. Cell Biol. 2, 531–539.

Joseph, B. B., Wang, Y., Edeen, P., Lažetić, V., Grant, B. D. and Fay, D. S.(2020). Control of clathrin-mediated endocytosis by NIMA family kinases. PLoS Genet. 16, e1008633.

Kamath, R. S. and Ahringer, J.(2003). Genome-wide RNAi screening in Caenorhabditis elegans. Methods San Diego Calif 30, 313–321.

Kodani, A., Tonthat, V., Wu, B. and Sütterlin, C.(2010). Par6α Interacts with the Dynactin Subunit p150Glued and Is a Critical Regulator of Centrosomal Protein Recruitment. Mol. Biol. Cell 21, 3376–3385.

Koorman, T., Klompstra, D., van der Voet, M., Lemmens, I., Ramalho, J. J., Nieuwenhuize, S., van den Heuvel, S., Tavernier, J., Nance, J. and Boxem, M.(2016). A combined binary interaction and phenotypic map of *C. elegans* cell polarity proteins. Nat. Cell Biol. 18, 337–346.

Krahn, M. P., Bückers, J., Kastrup, L. and Wodarz, A.(2010). Formation of a Bazooka-Stardust complex is essential for plasma membrane polarity in epithelia. J. Cell Biol. 190, 751–760.

Lažetić, V. and Fay, D. S.(2017). Molting in C. elegans. Worm 6, e1330246.

Leibfried, A., Fricke, R., Morgan, M. J., Bogdan, S. and Bellaiche, Y.(2008). Drosophila Cip4 and WASp define a branch of the Cdc42-Par6-aPKC pathway regulating E-cadherin endocytosis. Curr. Biol. CB 18, 1639–1648.

Lenfant, N., Polanowska, J., Bamps, S., Omi, S., Borg, J.-P. and Reboul, J.(2010). A genome-wide study of PDZ-domain interactions in C. elegans reveals a high frequency of non-canonical binding. BMC Genomics 11, 671.

Leung, B., Hermann, G. J. and Priess, J. R.(1999). Organogenesis of the Caenorhabditis elegans intestine. Dev. Biol. 216, 114–134.

Li, J., Kim, H., Aceto, D. G., Hung, J., Aono, S. and Kemphues, K. J.(2010a). Binding to PKC-3, but not to PAR-3 or to a conventional PDZ domain ligand, is required for PAR-6 function in C. elegans. Dev. Biol. 340, 88–98.

Li, B., Kim, H., Beers, M. and Kemphues, K.(2010b). Different domains of C. elegans PAR-3 are required at different times in development. Dev. Biol. 344, 745–757.

Lin, D., Edwards, A. S., Fawcett, J. P., Mbamalu, G., Scott, J. D. and Pawson, T.(2000). A mammalian PAR-3-PAR-6 complex implicated in Cdc42/Rac1 and aPKC signalling and cell polarity. Nat. Cell Biol. 2, 540–547.

Liu, X.-F., Ishida, H., Raziuddin, R. and Miki, T.(2004). Nucleotide exchange factor ECT2 interacts with the polarity protein complex Par6/Par3/protein kinase Czeta (PKCzeta) and regulates PKCzeta activity. Mol. Cell. Biol. 24, 6665–6675.

Liu, X. F., Ohno, S. and Miki, T.(2006). Nucleotide exchange factor ECT2 regulates epithelial cell polarity. Cell. Signal. 18, 1604–1615.

Luck, K., Kim, D.-K., Lambourne, L., Spirohn, K., Begg, B. E., Bian, W., Brignall, R., Cafarelli, T., Campos-Laborie, F. J., Charloteaux, B., et al.(2020). A reference map of the human binary protein interactome. Nature 580, 402–408.

Martin-Belmonte, F., Gassama, A., Datta, A., Yu, W., Rescher, U., Gerke, V. and Mostov, K.(2007). PTEN-mediated apical segregation of phosphoinositides controls epithelial morphogenesis through Cdc42. Cell 128, 383–397.

McCaffrey, L. M. and Macara, I. G.(2009). Widely conserved signaling pathways in the establishment of cell polarity. Cold Spring Harb. Perspect. Biol. 1, a001370.

McMahon, L., Legouis, R., Vonesch, J. L. and Labouesse, M.(2001). Assembly of C. elegans apical junctions involves positioning and compaction by LET-413 and protein aggregation by the MAGUK protein DLG-1. J. Cell Sci. 114, 2265–2277.

Meli, V. S., Osuna, B., Ruvkun, G. and Frand, A. R.(2010). MLT-10 Defines a Family of DUF644 and Proline-rich Repeat Proteins Involved in the Molting Cycle of Caenorhabditis elegans. Mol. Biol. Cell 21, 1648–1661.

Mertens, A. E. E., Rygiel, T. P., Olivo, C., van der Kammen, R. and Collard, J. G.(2005). The Rac activator Tiam1 controls tight junction biogenesis in keratinocytes through binding to and activation of the Par polarity complex. J. Cell Biol. 170, 1029–1037.

Montoyo-Rosario, J. G., Armenti, S. T., Zilberman, Y. and Nance, J.(2020). The Role of pkc-3 and Genetic Suppressors in Caenorhabditis elegans Epithelial Cell Junction Formation. Genetics 214, 941–959.

Morais-de-Sá, E., Mirouse, V. and St Johnston, D.(2010). aPKC phosphorylation of Bazooka defines the apical/lateral border in Drosophila epithelial cells. Cell 141, 509–523.

Motegi, F., Zonies, S., Hao, Y., Cuenca, A. A., Griffin, E. and Seydoux, G.(2011). Microtubules induce self-organization of polarized PAR domains in Caenorhabditis elegans zygotes. Nat. Cell Biol. 13, 1361–1367.

Myers, T. R. and Greenwald, I.(2005). lin-35 Rb Acts in the Major Hypodermis to Oppose Ras-Mediated Vulval Induction in C. elegans. Dev. Cell 8, 117–123.

Nagai-Tamai, Y., Mizuno, K., Hirose, T., Suzuki, A. and Ohno, S.(2002). Regulated protein-protein interaction between aPKC and PAR-3 plays an essential role in the polarization of epithelial cells. Genes Cells Devoted Mol. Cell. Mech. 7, 1161–1171.

Nance, J. and Priess, J. R.(2002). Cell polarity and gastrulation in C. elegans. Dev. Camb. Engl. 129, 387–397.

Nance, J., Munro, E. M. and Priess, J. R.(2003). C. elegans PAR-3 and PAR-6 are required for apicobasal asymmetries associated with cell adhesion and gastrulation. Dev. Camb. Engl. 130, 5339–5350.

Nunes de Almeida, F., Walther, R. F., Pressé, M. T., Vlassaks, E. and Pichaud, F.(2019). Cdc42 defines apical identity and regulates epithelial morphogenesis by promoting apical recruitment of Par6-aPKC and Crumbs. Dev. Camb. Engl. 146,.

Ozdamar, B., Bose, R., Barrios-Rodiles, M., Wang, H.-R., Zhang, Y. and Wrana, J. L.(2005). Regulation of the polarity protein Par6 by TGFbeta receptors controls epithelial cell plasticity. Science 307, 1603–1609.

Petronczki, M. and Knoblich, J. A.(2001). DmPAR-6 directs epithelial polarity and asymmetric cell division of neuroblasts in Drosophila. Nat. Cell Biol. 3, 43–49.

Pickett, M. A., Naturale, V. F. and Feldman, J. L.(2019). A Polarizing Issue: Diversity in the Mechanisms Underlying Apico-Basolateral Polarization In Vivo. Annu. Rev. Cell Dev. Biol. 35, 285–308.

Pinal, N., Goberdhan, D. C. I., Collinson, L., Fujita, Y., Cox, I. M., Wilson, C. and Pichaud, F.(2006). Regulated and polarized PtdIns(3,4,5)P3 accumulation is essential for apical membrane morphogenesis in photoreceptor epithelial cells. Curr. Biol. CB 16, 140–149.

Plant, P. J., Fawcett, J. P., Lin, D. C. C., Holdorf, A. D., Binns, K., Kulkarni, S. and Pawson, T.(2003). A polarity complex of mPar-6 and atypical PKC binds, phosphorylates and regulates mammalian Lgl. Nat. Cell Biol. 5, 301–308.

Ramalho, J. J., Sepers, J. J., Nicolle, O., Schmidt, R., Cravo, J., Michaux, G. and Boxem, M.(2020). C-terminal phosphorylation modulates ERM-1 localization and dynamics to control cortical actin organization and support lumen formation during C. elegans development. Dev. Camb. Engl.

Ramanujam, R., Han, Z., Zhang, Z., Kanchanawong, P. and Motegi, F.(2018). Establishment of the PAR-1 cortical gradient by the aPKC-PRBH circuit. Nat. Chem. Biol. 14, 917–927.

Rodriguez, J., Peglion, F., Martin, J., Hubatsch, L., Reich, J., Hirani, N., Gubieda, A. G., Roffey, J., Fernandes, A. R., St Johnston, D., et al.(2017). aPKC Cycles between Functionally Distinct PAR Protein Assemblies to Drive Cell Polarity. Dev. Cell 42, 400–415.e9.

Rodriguez-Boulan, E. and Macara, I. G.(2014). Organization and execution of the epithelial polarity programme. Nat. Rev. Mol. Cell Biol. 15, 225–242.

Rosa, A., Vlassaks, E., Pichaud, F. and Baum, B.(2015). Ect2/Pbl acts via Rho and polarity proteins to direct the assembly of an isotropic actomyosin cortex upon mitotic entry. Dev. Cell 32, 604–616.

Rual, J.-F., Ceron, J., Koreth, J., Hao, T., Nicot, A.-S., Hirozane-Kishikawa, T., Vandenhaute, J., Orkin, S. H., Hill, D. E., van den Heuvel, S., et al.(2004). Toward Improving Caenorhabditis elegans Phenome Mapping With an ORFeome-Based RNAi Library. Genome Res. 14, 2162–2168.

Rueden, C. T., Schindelin, J., Hiner, M. C., DeZonia, B. E., Walter, A. E., Arena, E. T. and Eliceiri, K. W.(2017). ImageJ2: ImageJ for the next generation of scientific image data. BMC Bioinformatics 18, 529.

Russel, S., Frand, A. R. and Ruvkun, G.(2011). Regulation of the C. elegans molt by pqn-47. Dev. Biol. 360, 297–309.

Sallee, M. D., Zonka, J. C., Skokan, T. D., Raftrey, B. C. and Feldman, J. L.(2018). Tissue-specific degradation of essential centrosome components reveals distinct microtubule populations at microtubule organizing centers. PLOS Biol. 16, e2005189.

Sánchez, N. S. and Barnett, J. V.(2012). TGFβ and BMP-2 regulate epicardial cell invasion via TGFβR3 activation of the Par6/Smurf1/RhoA pathway. Cell. Signal. 24, 539–548.

Sanchez, A. D. and Feldman, J. L.(2017). Microtubule-organizing centers: from the centrosome to non-centrosomal sites. Curr. Opin. Cell Biol. 44, 93–101.

Schiestl, R. H. and Gietz, R. D.(1989). High efficiency transformation of intact yeast cells using single stranded nucleic acids as a carrier. Curr. Genet. 16, 339–346.

Schindelin, J., Arganda-Carreras, I., Frise, E., Kaynig, V., Longair, M., Pietzsch, T., Preibisch, S., Rueden, C., Saalfeld, S., Schmid, B., et al.(2012). Fiji: an open-source platform for biological-image analysis. Nat. Methods 9, 676–682.

Schober, M., Schaefer, M. and Knoblich, J. A.(1999). Bazooka recruits Inscuteable to orient asymmetric cell divisions in Drosophila neuroblasts. Nature 402, 548–551.

Schwartz, M. L. and Jorgensen, E. M.(2016). SapTrap, a Toolkit for High-Throughput CRISPR/Cas9 Gene Modification in *Caenorhabditis elegans*. Genetics 202, 1277–1288.

Shafaq-Zadah, M., Brocard, L., Solari, F. and Michaux, G.(2012). AP-1 is required for the maintenance of apico-basal polarity in the C. elegans intestine. Development 139, 2061–2070.

Shahab, J., Tiwari, M. D., Honemann-Capito, M., Krahn, M. P. and Wodarz, A.(2015). Bazooka/PAR3 is dispensable for polarity in Drosophila follicular epithelial cells. Biol. Open 4, 528–541.

St Johnston, D.(2018). Establishing and transducing cell polarity: common themes and variations. Curr. Opin. Cell Biol. 51, 33–41.

Tabuse, Y., Izumi, Y., Piano, F., Kemphues, K. J., Miwa, J. and Ohno, S.(1998). Atypical protein kinase C cooperates with PAR-3 to establish embryonic polarity in Caenorhabditis elegans. Dev. Camb. Engl. 125, 3607–3614.

Taffoni, C., Omi, S., Huber, C., Mailfert, S., Fallet, M., Rupprecht, J.-F., Ewbank, J. J. and Pujol, N.(2020). Microtubule plus-end dynamics link wound repair to the innate immune response. eLife 9,.

Tinevez, J.-Y., Perry, N., Schindelin, J., Hoopes, G. M., Reynolds, G. D., Laplantine, E., Bednarek, S. Y., Shorte, S. L. and Eliceiri, K. W.(2017). TrackMate: An open and extensible platform for single-particle tracking. Methods 115, 80–90.

Totong, R., Achilleos, A. and Nance, J.(2007). PAR-6 is required for junction formation but not apicobasal polarization in C. elegans embryonic epithelial cells. Dev. Camb. Engl. 134, 1259–1268.

von Stein, W., Ramrath, A., Grimm, A., Müller-Borg, M. and Wodarz, A.(2005). Direct association of Bazooka/PAR-3 with the lipid phosphatase PTEN reveals a link between the PAR/aPKC complex and phosphoinositide signaling. Development 132, 1675–1686.

Von Stetina, S. E. and Mango, S. E.(2015). PAR-6, but not E-cadherin and β-integrin, is necessary for epithelial polarization in C. elegans. Dev. Biol. 403, 5–14.

Von Stetina, S. E., Liang, J., Marnellos, G. and Mango, S. E.(2017). Temporal regulation of epithelium formation mediated by FoxA, MKLP1, MgcRacGAP, and PAR-6. Mol. Biol. Cell 28, 2042–2065.

Waaijers, S., Ramalho, J. J., Koorman, T., Kruse, E. and Boxem, M.(2015). The C. elegans Crumbs family contains a CRB3 homolog and is not essential for viability. Biol. Open 4, 276–284.

Waaijers, S., Muñoz, J., Berends, C., Ramalho, J. J., Goerdayal, S. S., Low, T. Y., Zoumaro-Djayoon, A. D., Hoffmann, M., Koorman, T., Tas, R. P., et al.(2016). A tissue-specific protein purification approach in Caenorhabditis elegans identifies novel interaction partners of DLG-1/Discs large. BMC Biol. 14,.

Walther, R. F. and Pichaud, F.(2010). Crumbs/DaPKC-dependent apical exclusion of Bazooka promotes photoreceptor polarity remodeling. Curr Biol 20, 1065–74.

Wang, H.-R., Zhang, Y., Ozdamar, B., Ogunjimi, A. A., Alexandrova, E., Thomsen, G. H. and Wrana, J. L.(2003). Regulation of cell polarity and protrusion formation by targeting RhoA for degradation. Science 302, 1775–1779.

Wang, S., Wu, D., Quintin, S., Green, R. A., Cheerambathur, D. K., Ochoa, S. D., Desai, A. and Oegema, K.(2015). NOCA-1 functions with γ-tubulin and in parallel to Patronin to assemble non-centrosomal microtubule arrays in C. elegans. eLife 4, e08649.

Wang, S.-C., Low, T. Y. F., Nishimura, Y., Gole, L., Yu, W. and Motegi, F.(2017). Cortical forces and CDC-42 control clustering of PAR proteins for Caenorhabditis elegans embryonic polarization. Nat. Cell Biol. 19, 988–995.

Watts, J. L., Etemad-Moghadam, B., Guo, S., Boyd, L., Draper, B. W., Mello, C. C., Priess, J. R. and Kemphues, K. J.(1996). par-6, a gene involved in the establishment of asymmetry in early C. elegans embryos, mediates the asymmetric localization of PAR-3. Dev. Camb. Engl. 122, 3133–3140.

Welchman, D. P., Mathies, L. D. and Ahringer, J.(2007). Similar requirements for CDC-42 and the PAR-3/PAR-6/PKC-3 complex in diverse cell types. Dev. Biol. 305, 347–357.

Wildwater, M., Sander, N., de Vreede, G. and van den Heuvel, S.(2011). Cell shape and Wnt signaling redundantly control the division axis of *C. elegans* epithelial stem cells. Development 138, 4375–4385.

Wilson, M. I., Gill, D. J., Perisic, O., Quinn, M. T. and Williams, R. L.(2003). PB1 domain-mediated heterodimerization in NADPH oxidase and signaling complexes of atypical protein kinase C with Par6 and p62. Mol. Cell 12, 39–50.

Winter, J. F., Höpfner, S., Korn, K., Farnung, B. O., Bradshaw, C. R., Marsico, G., Volkmer, M., Habermann, B. and Zerial, M.(2012). *Caenorhabditis elegans* screen reveals role of PAR-5 in RAB-11-recycling endosome positioning and apicobasal cell polarity. Nat. Cell Biol. 14, 666–676.

Wodarz, A., Ramrath, A., Kuchinke, U. and Knust, E.(1999). Bazooka provides an apical cue for Inscuteable localization in Drosophila neuroblasts. Nature 402, 544–547.

Wodarz, A., Ramrath, A., Grimm, A. and Knust, E.(2000). Drosophila atypical protein kinase C associates with Bazooka and controls polarity of epithelia and neuroblasts. J. Cell Biol. 150, 1361–1374.

Yamanaka, T., Horikoshi, Y., Suzuki, A., Sugiyama, Y., Kitamura, K., Maniwa, R., Nagai, Y., Yamashita, A., Hirose, T., Ishikawa, H., et al.(2001). PAR-6 regulates aPKC activity in a novel way and mediates cell-cell contact-induced formation of the epithelial junctional complex. Genes Cells Devoted Mol. Cell. Mech. 6, 721–731.

Yamanaka, T., Horikoshi, Y., Sugiyama, Y., Ishiyama, C., Suzuki, A., Hirose, T., Iwamatsu, A., Shinohara, A. and Ohno, S.(2003). Mammalian Lgl forms a protein complex with PAR-6 and aPKC independently of PAR-3 to regulate epithelial cell polarity. Curr. Biol. CB 13, 734–743.

Yochem, J., Tuck, S., Greenwald, I. and Han, M.(1999). A gp330/megalin-related protein is required in the major epidermis of Caenorhabditis elegans for completion of molting. Dev. Camb. Engl. 126, 597–606.

Yochem, J., Lažetić, V., Bell, L., Chen, L. and Fay, D.(2015). C. elegans NIMA-related kinases NEKL-2 and NEKL-3 are required for the completion of molting. Dev. Biol. 398, 255–266.

Yu, H., Braun, P., Yildirim, M. A., Lemmens, I., Venkatesan, K., Sahalie, J., Hirozane-Kishikawa, T., Gebreab, F., Li, N., Simonis, N., et al.(2008). High-quality binary protein interaction map of the yeast interactome network. Science 322, 104–110.

Zhang, H., Abraham, N., Khan, L. A., Hall, D. H., Fleming, J. T. and Göbel, V.(2011). Apicobasal domain identities of expanding tubular membranes depend on glycosphingolipid biosynthesis. Nat. Cell Biol. 13, 1189–1201.

Zhang, H., Kim, A., Abraham, N., Khan, L. A., Hall, D. H., Fleming, J. T. and Gobel, V.(2012). Clathrin and AP-1 regulate apical polarity and lumen formation during C. elegans tubulogenesis. Dev. Camb. Engl. 139, 2071–2083.

Zhang, L., Ward, J. D., Cheng, Z. and Dernburg, A. F.(2015). The auxin-inducible degradation (AID) system enables versatile conditional protein depletion in C. elegans. Dev. Camb. Engl. 142, 4374–4384.

