## Supplemental Figures for "Epidermal PAR-6 and PKC-3 are essential for postembryonic development of *Caenorhabditis elegans* and control non-centrosomal microtubule organization"

### Figure S1

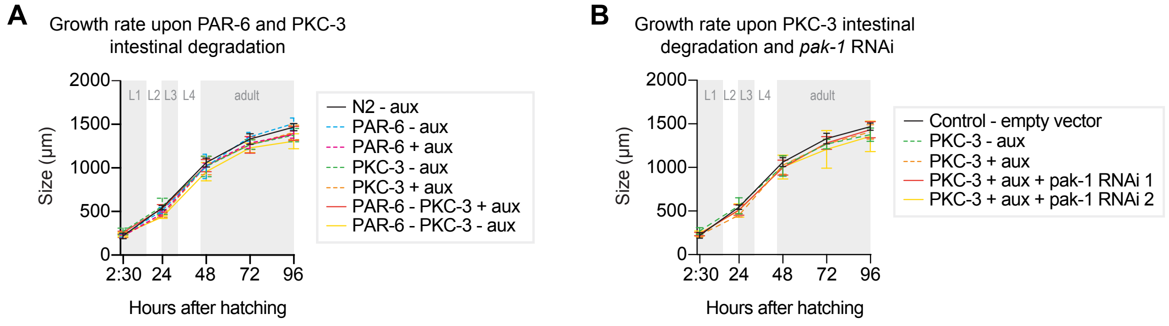


**Fig. S1. PKC-3 does not act redundantly with PAR-6 or PAK-1 in the *C. elegans* intestine. (A)** Growth curves of N2, *par-6::aid::gfp, gfp::aid::pkc-3*, and *par-6::aid::gfp; gfp::aid::pkc-3* (double depletion) animals in absence (- aux) or presence (+ aux) of 4 mM auxin. Data of single PAR-6 and PKC-3 depletions (dashed lines) are repeated from Fig. 1 for comparison. Data show mean ± SD. *n* = 6, 7, 8, and 8 for N2 - aux; 6, 7, 9, and 9 for N2 + aux; 7, 6, 9, and 9 for PAR-6 - aux; 8, 6, 7, and 9 for PAR-6 + aux; 22, 11, 10, and 14 for PKC-3 - aux; 19, 14, 9, and 10 for PKC-3 + aux; 5, 8, 6, 8, and 7 for PAR-6–PKC-3 - aux, and 11, 7, 10, 7, and 8 for PAR-6–PKC-3 + aux. **(B)** Growth curves of N2 and *gfp::aid::pkc-3* animals in absence (- aux) or presence (+ aux) of 4 mM auxin and upon feeding of *pak-1* RNAi. *pak-1* RNAi clones were obtained from the Ahringer (RNAi 1) or Vidal (RNAi 2) genome-wide RNAi libraries. Data of single PKC-3 depletion (dashed lines) is repeated from Fig. 1 for comparison. Data show mean ± SD. *n* = 13, 10, 13, 14, and 12 for Control – empty vector; 8, 7, 8, 4, and 9 for PKC-3 – aux; 8, 7, 8, 8, and 8 for PKC-3 + aux; 10, 13, 18, 10, and 15 for PKC-3 + aux + pak-1 RNAi 1 and 14, 7, 14, 13, and 7 for PKC-3 + aux + pak-1 RNAi 2.

### Figure S2

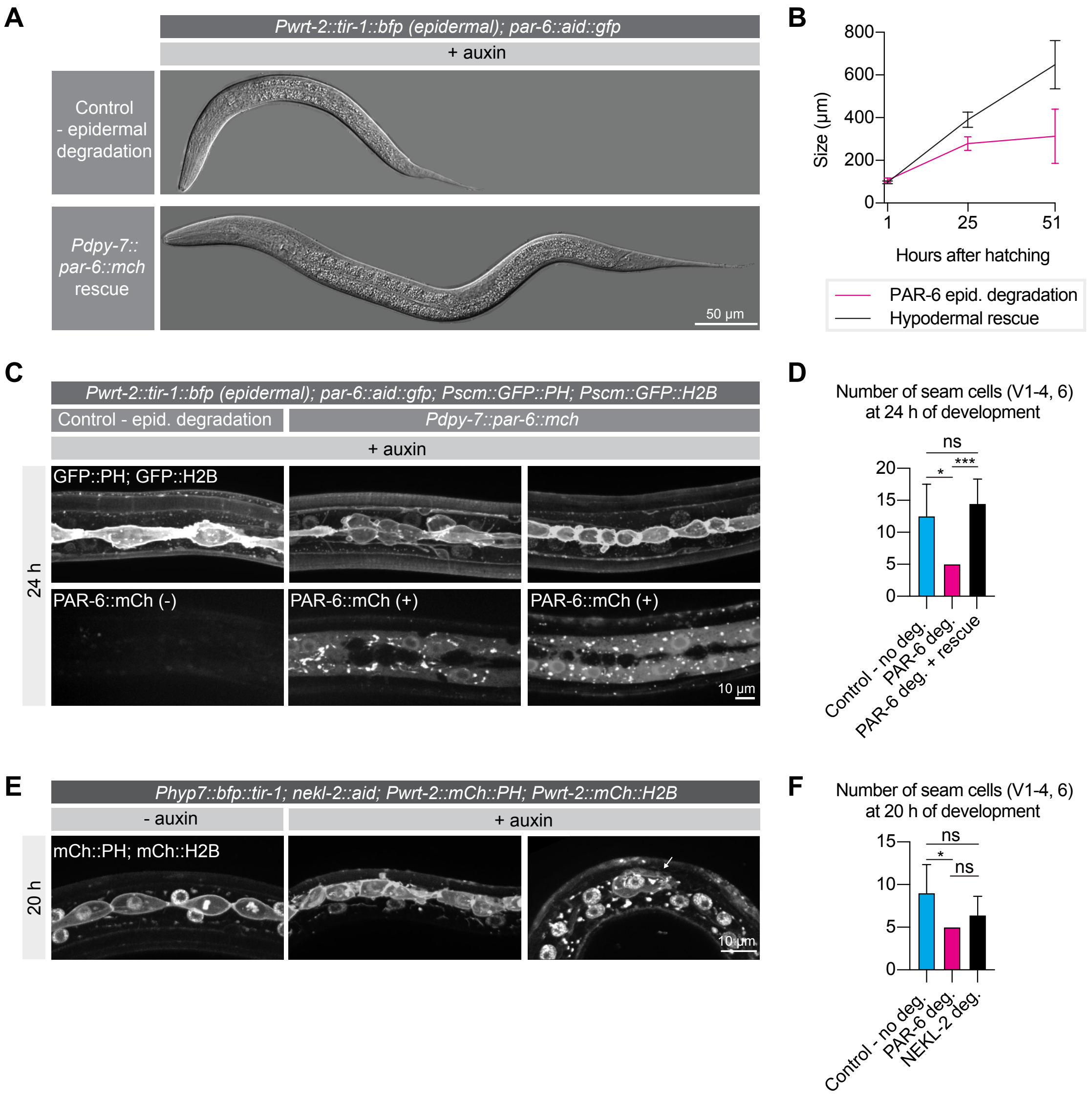


**Fig. S2. Hypodermal expression of PAR-6 is necessary for larval development. (A)** DIC microscopy images of *par-6::aid::gfp; Pwrt-2::tir-1::bfp* animals carrying (*Pdpy-7::par-6::mCherry*) or lacking (Control) an extrachromosomal array expressing PAR-6 in the hypodermis. Worms were grown in presence (+ auxin) of 4 mM auxin since hatching. Images taken 30 h after hatching. **(B)** Growth curves of animals treated as in (A). Lengths were measured at 1, 25, and 51 h after hatching. *n* = 7, 8, and 4 for PAR-6 epidermal degradation and 11, 14, and 14 for hypodermal rescue. Data shows mean ± SD. **(C)** Seam cells visualized by nuclear H2B::GFP and membrane-bound PH::GFP at 24 h of post-embryonic development in *par-6::aid::gfp* animals carrying (*Pdpy-7::par-6::mCherry*) or lacking (Control) an extrachromosomal array expressing PAR-6 in the hypodermis. Worms were grown in presence (+ auxin) of 4 mM auxin since hatching. **(D)** Number of seam cells (V1–4, 6) at 24 h of post-embryonic development in absence (- auxin) or presence (+ auxin) of 4 mM auxin and with or without PAR-6 hypodermal rescue. *n* = 4 for control, 5 for PAR-6 deg, and 11 for PAR-6 deg + rescue. Bars represent mean ± SD. **(E)** Seam cells visualized by nuclear H2B::mCherry and membrane-bound PH::mCherry at 20 h of post-embryonic development in *nekl-2::aid; Phyp7::bfp::tir-1* animals in absence (- aux) or presence (+ aux) of 4 mM auxin since hatching. **(F)** Number of seam cells (V1–4, 6) at 20 h of post-embryonic development in *nekl-2::aid; Phyp7::bfp::tir-1* animals in absence (Control – no deg) or presence (NEKL-2 deg.) of 4 mM auxin and in *par-6::aid::gfp; Pwrt-2::tir-1::bfp* animals in presence (PAR-6 deg.) of 4 mM auxin. *n* = 6 for Control, 5 for PAR-6 deg, and 10 for NEKL-2 deg. Bars represent mean ± SD. Tests of significance: Tukey's test of significance for D and F. ns = not significant, * = P ≤ 0.05, *** = P ≤ 0.001.

### Figure S3

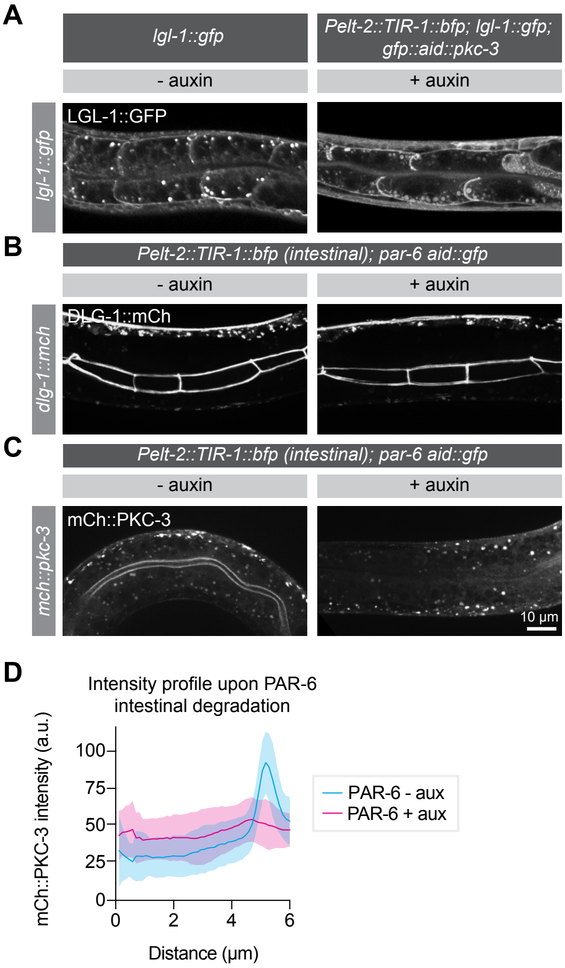


**Fig. S3. PAR-6 and PKC-3 are not essential for LGL-1 localization or junction maintenance in the larval intestine.** **(A)** Distribution of LGL-1::GFP, in *lgl-1::gfp* animals without auxin and in *lgl-1::gfp; gfp::aid::pkc-3; Pelt-2::tir-1::bfp* animals in presence of 4 mM auxin. Images are maximum projections covering the whole intestinal cells. **(B)** Distribution of DLG-1::mCherry in *par-6::aid::gfp; Pelt-2::tir-1::bfp* animals in absence (- auxin) or presence (+ auxin) of 4 mM auxin. Images are maximum projections of the luminal domain for the intestine. **(C, D)** Distribution and quantification of mCherry::PKC-3 in *pkc-3::mCherry; par-6::aid::gfp; Pelt-2::tir-1::bfp* animals in absence (- aux) or presence (+ aux) of 1 mM auxin. Images are maximum projections of the apical domain. Quantifications shows mean apical GFP fluorescence intensity ± SD at the hyp7–seam cell junction. n = for 8 PAR-6 - aux and 4 for PAR-6 + aux.

### Figure S4

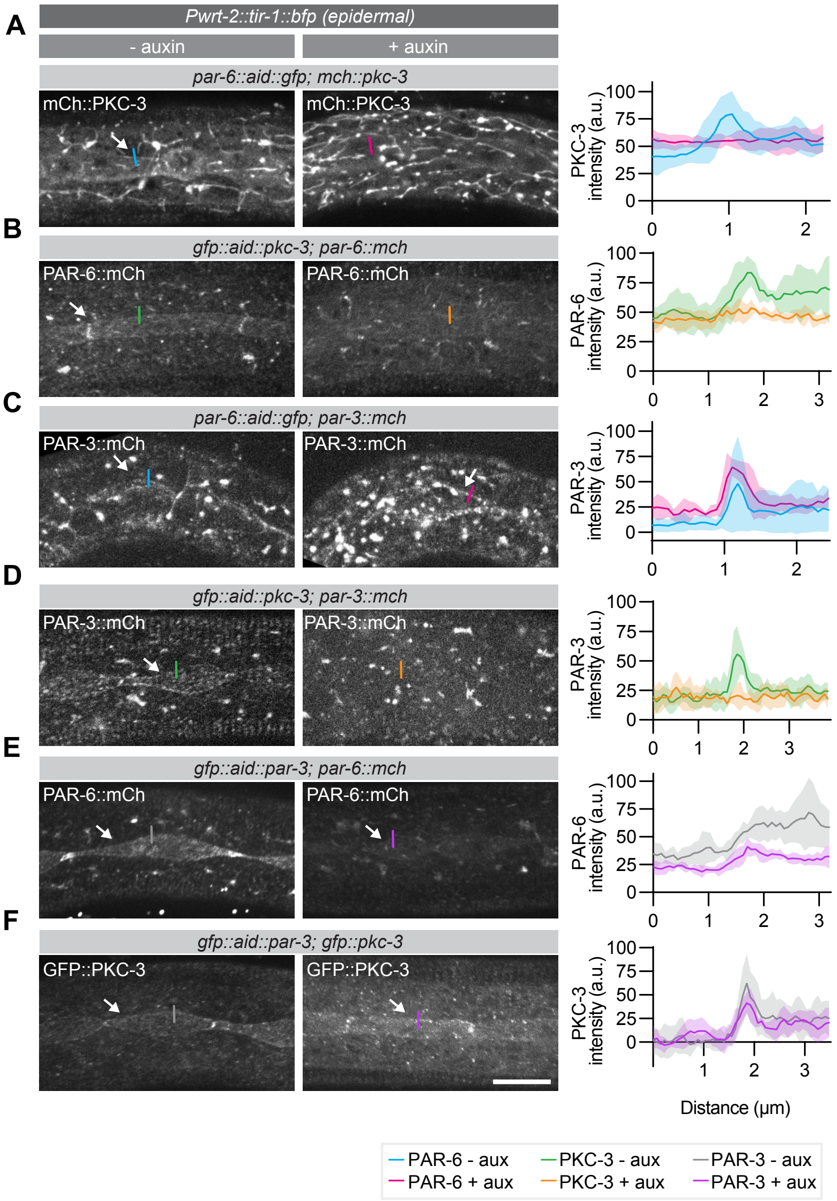


**Fig. S4. Localization dependencies of PAR-6, PKC-3, and PAR-3 in the seam cells.** Left panels are of animals not exposed to auxin, right panels are of animals exposed to 1 mM auxin for 1 h. **(A)** Distribution of mCherry::PKC-3 in *par-6::aid::gfp* animals. **(B)** Distribution of PAR-6::mCherry in *gfp::aid::pkc-3* animals. **(C, D)** Distribution of PAR-3::mCherry in *par-6::aid::gfp* and *gfp::aid::pkc-3* animals. **(E, F)** PAR-6::mCherry and GFP::PKC-3 localization *in gfp::aid::par-3* animals. Images are maximum projections of a z-stack. Quantifications show mean apical GFP fluorescence intensity ± SD at the hyp7–seam cell junction. n = 5 animals for all data points.

### Figure S5

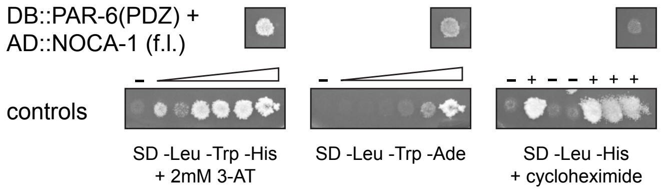


**Fig. S5. Interaction of PAR-6 and NOCA-1 in the yeast two hybrid system.** The PAR-6 PDZ domain fused to the Gal4 DNA binding domain was co-expressed with full-length NOCA-1 fused to the Gal4 activation domain. Growth on -Leu -Trp -His + 2mM 3-AT, and on -Leu -Trp -Ade plates indicates presence of interaction. Lack of growth on -Leu -His + cycloheximide plate shows that DB::PAR-6 is not self-activating. Controls range from no reporter activation to strong reporter activation. On cycloheximide plates, (-) indicates no growth expected, and (+) indicates growth expected.
